# Predicting anti-cancer drug synergy using extended drug similarity profiles

**DOI:** 10.1101/2022.08.28.505568

**Authors:** Sayed-Rzgar Hosseini, Xiaobo Zhou

## Abstract

Combination therapy is a promising strategy for confronting the complexity of cancer. However, experimental exploration of the vast space of potential drug combinations is costly and unfeasible. Therefore, computational methods for predicting drug synergy are much-needed for narrowing down this space, especially when examining new cellular contexts. Here, we thus introduce CCSynergy, a flexible, context-aware and integrative deep learning framework that we have established to unleash the potential of the Chemical Checker extended drug similarity profiles for the purpose of drug synergy prediction. We have shown that CCSynergy enables predictions of superior accuracy, remarkable robustness and improved context-generalizability as compared to the state-of-the-art methods in the field. Having established the potential of CCSynergy for generating experimentally validated predictions, we exhaustively explored the untested drug combination space. This resulted in a compendium of potentially synergistic drug combinations on hundreds of cancer cell lines, which can guide future experimental screens.

## Introduction

Aberrant behavior of cancer cells is caused by malfunctioning of multiple signaling pathways that promote proliferation and inhibit apoptosis^1^. The pervasive redundancy, inherent multifunctionality and the combinatorial control of these biological processes, have challenged the traditional “one gene, one drug” paradigm pioneered by Ehrlich^2,3^. This is evidenced by the increasing rate of drug failure and the recurrent emergence of drug resistance in targeted cancer therapy^2,4^. To overcome these challenges, combination therapy is a promising strategy as drug synergy ensures greater efficacy in lower drug dosages, which results in avoiding toxicity and minimizing the chance of drug resistance^5^.

High-throughput screening methods have enabled testing and quantifying drug synergy^6^. However, synergistic drug pairs are rare and exhaustive exploration of the vast space of potential drug combinations is not experimentally feasible. Thus, computational predictive models of drug synergy, which enable prioritization of the candidate drug combinations, are much-needed for narrowing down this vast search space. Computational models as diverse as kinetic^7,8^, network^9,10^ and logic models^11,12^ have thus been employed to gain quantitative insights into the mystery of drug synergy. Moreover, the availability of large-scale drug synergy data, such as the Merck^13^ dataset, has encouraged the emergence of a wide variety of machine learning based methods ranging from logistic regression^14^ to extremely randomized trees^15^ and XGBoost^16^. Ultimately, the state-of-the-art deep learning approaches such as DeepSynergy^17^ and recently TranSynergy^18^ have entered the race and outperformed the others.

Various metrics of drug similarity have been proposed to represent drug pairs in drug synergy prediction models. The focus has primarily been on chemical features of drugs^17,19–21^, and structural or network-level similarity of their targets within cells^9,22–24^. Further quantities based on phenotypic effects of drugs such as therapeutic and side effect similarities or cell-line based sensitivity profiles have also been considered^25–27^. Moreover, the Connectivity Map^28^ has sparked development of new similarity metrics based on drug-induced gene expression profiles^29–33^. Not surprisingly, different combinations of these similarity measures have also been examined^34–39^. Ultimately, the Chemical Checker (CC) database arose, which provides a unified framework to systematically extend the concept of drug similarity to all levels of biology, from chemistry, targets, networks, to cellular and clinical effects of drugs^40^. Nevertheless, its enormous potential for predicting drug synergy has not yet been unleashed. This motivated us to take the pioneering step to develop a new drug synergy prediction framework by integrating all 25 levels of CC bioactivity similarity metrics (**Fig. 1a**).

**Figure 1.**
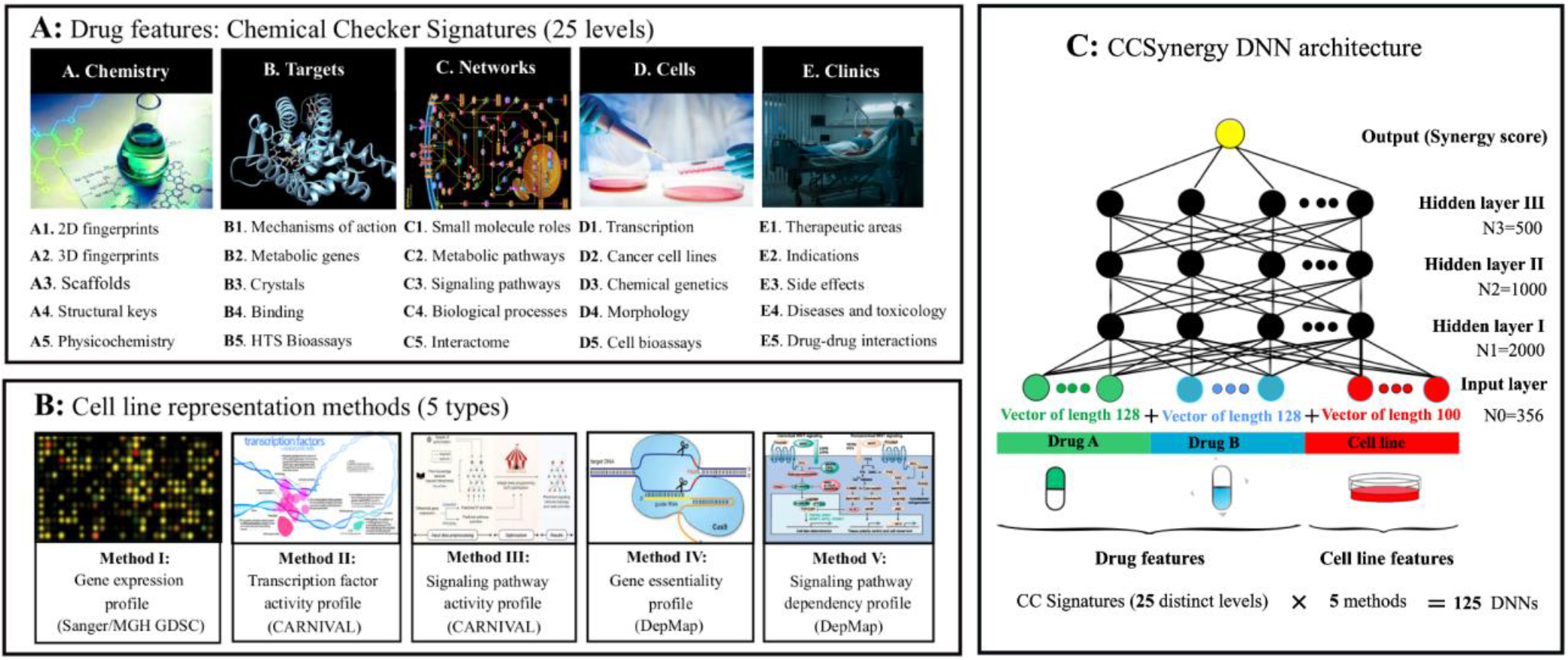
CCSynergy framework. **A)** The Chemical Checker^40^ based signatures of type II are used as drug features in CCSynergy, which cover five main characteristics of small molecules: A. chemistry, B. targets, C. networks, D. cells and E. clinics. Each of them is further divided into five sub-categories totaling 25 distinct levels for drug representation. **B)** Five different methods for representing cancer cell lines are used: I) down-stream gene expression profiles, II) transcription factor activity and III) signaling pathway activity profiles inferred using CARNIVAL^46^, IV) Dep-Map based gene essentiality and V) signaling pathway dependency profiles (see **Methods**). **C)** CCSynergy DNN architecture. A given triplet is represented as a vector of length 356, which is formed by concatenating the corresponding vectors of drug pair and cell line features. Since drug synergy is order-agnostic, each triplet is represented twice to account for both directions (AB and BA). The DNN contains three hidden layers comprising 2000, 1000 and 500 neurons respectively, which propagate information from the input layer to the output unit. 125 distinct DNNs are trained, each corresponding to one of the 25 CC spaces and one of the 5 cell line representation methods.

Moreover, it is well-established that drug synergy is highly context specific^41^, which necessitates precise representation of the cellular features in drug synergy prediction models. This is important, especially in precision medicine, where computational methods are required to enable accurate predictions in specific cellular contexts. Genome-wide expression profiles of cancer cell lines have been extensively employed for this purpose^17,18,20,23,42–45^. However, the expression level of downstream genes is not necessarily a strong indicator of the functional status of the cell and may not directly connect with the drug response phenotype. To address this, computational methods to infer the causal upstream processes, namely transcription factor or signaling pathway activities, which drive the downstream expression changes, have been introduced (e.g., CARNIVAL^46^). However, their potential for representation of the cell in drug synergy prediction, has not yet been unlocked. Moreover, genome-wide CRISPR-based essentiality profile of cancer cell lines (DepMap)^47–50^, which can more directly establish causal links with cell survival, constitutes another promising cell representation alternative. Therefore, we aimed to systematically examine and compare these methods for representing cellular contexts (**Fig. 1b**).

In this work, we thus introduce CCSynergy, a Chemical-Checker harnessing deep neural network that enables context-aware anti-cancer drug synergy prediction (**Fig. 1c**). Using our rigorous cross validation schemes (**Fig. 2**), we ensure that CCSynergy offers drug synergy predictions of superior accuracy, remarkable robustness and improved context-generalizability. Moreover, leveraging a recently published large-scale resource of drug synergy^41^, we extensively examine the potential of CCSynergy to generate experimentally validated predictions. Finally, we provide a compendium of potentially synergistic drug combinations that calls for follow-up experimental investigation.

**Figure 2.**
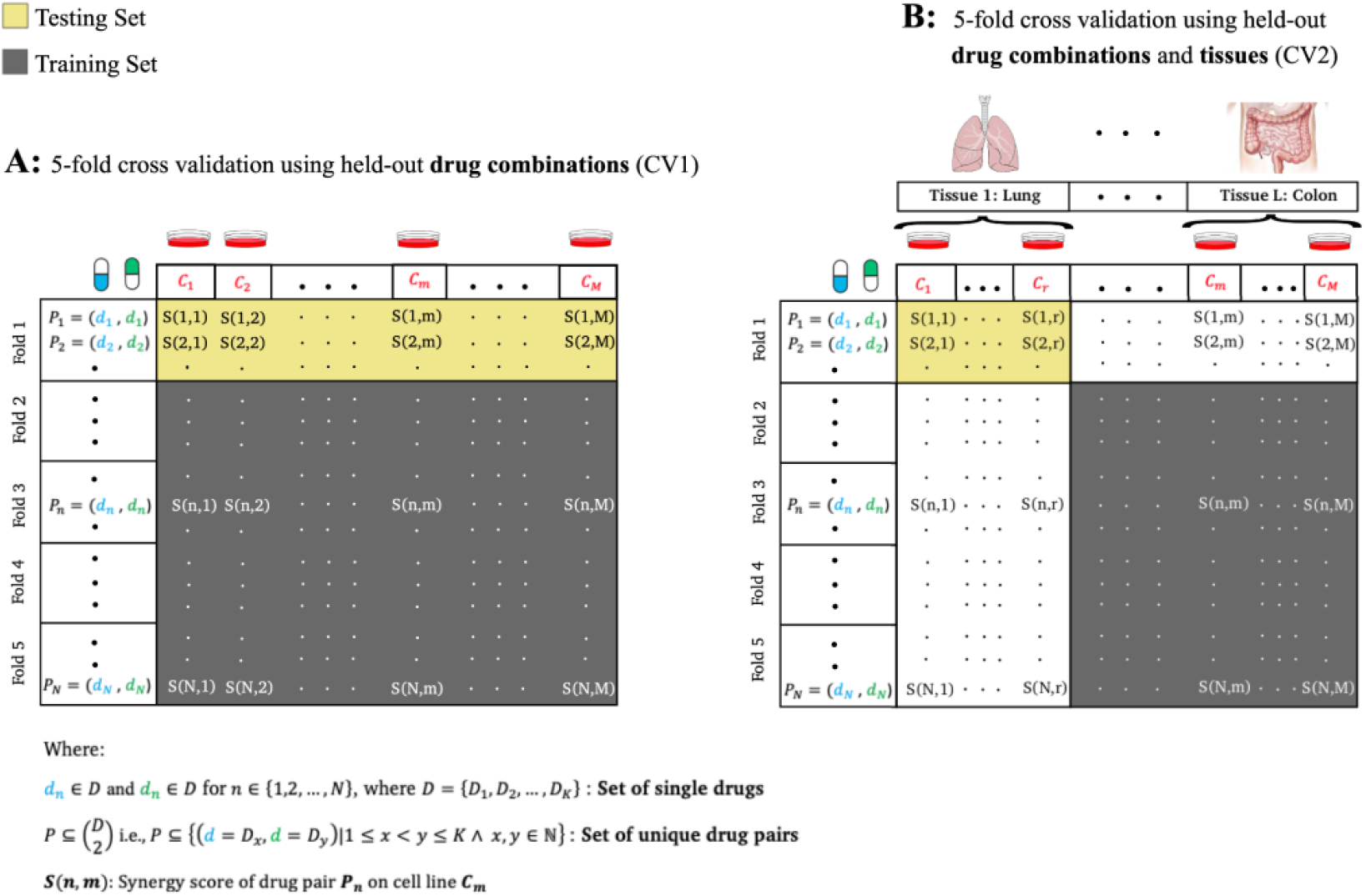
Cross validation schemes. A drug synergy dataset is shown as a matrix each element of which represents the synergy score *S*(*n, m*) of a given triplet of drug pair (*P*_*n*_) + cell line (*C*_*m*_). The dataset is divided row-wise into five-folds of equal size, which are needed in our 5-fold cross-validation schemes. **A)** CV1: 5 training cycles are needed in each of which, one fold is considered as the testing set (yellow) and the remaining four are used as the training set (gray). This ensures that the set of drug pairs in the testing and training sets do not overlap (held-out drug combinations). **B)** CV2: The matrix is not only divided row-wise, but also column-wise. Columns are grouped according to the tissue of origin of the corresponding cell lines. 5 × *L* learning cycles are needed, where L is the number of distinct tissues in our dataset. In this example, the elements corresponding to the rows in the first fold and a set of columns belonging to the lung tissue {*C*_1_,…,*C*_*r*_} are considered as the testing set (yellow). The samples in the remaining four folds (excluding those whose cell lines originated from the lung tissue), are considered as the training set (gray). This CV scheme further ensures that the cell lines in the training set originate from tissues that do not overlap with that of the testing set (held-out tissues).

## Results

### CCSynergy overview

The primary aim of CCSynergy is to unlock the potential of Chemical Checker (CC) bioactivity profiles^40^ for predicting anti-cancer drug synergy. CC catalogs integrated bioactivity data on almost 800,000 small molecules. It encompasses five levels of increasing complexity from A: the chemical properties of the compounds, B: their targets, and C: network-level properties, to D: their cellular, and E: clinical effects. Furthermore, each level is divided into five sub-levels resulting in 25 distinct signatures (**Fig. 1a**), each of which is represented in a vector format of the same length (128). The vectors have been generated via a two-step procedure: applying a dimensionality reduction technique (type I signatures) followed by running a network embedding approach on the resulting similarity networks (type II signatures. See **Methods** section).

CCSynergy also strives to enable context-aware predictions using five distinct methods for representing cellular contexts (**Fig. 1b**). I: the downstream gene expression profiles, II: the inferred transcription factor activity profiles, III: the inferred signaling pathway activity profiles, IV: CRISPR-based gene essentiality profiles, and V: DepMap based signaling pathway dependency profiles (See **Methods**). It is important to note that in CCSynergy, the cell lines are represented as vectors of the same lengths (100) after reducing the dimension of the original profiles either using auto-encoder based techniques (CCSynergy I and IV) or by selecting the top 100 most informative signaling pathways (CCSynergy III and V) or transcription factors (CCSynergy II).

Finally, we represent each sample as a vector of length 356 by concatenating its drug pair and cell line vectors (**Fig. 1c**). CCSynergy is a feed-forward deep neural network (DNN) and its architecture includes three hidden layers, which propagate information from the input vectors to the output unit, where the synergy score is predicted. For a given training set, 125 separate DNNs are trained each corresponding to one of the 25 CC spaces and one of the 5 cell line representation methods. Note that we considered different hyper-parameter settings and found the optimal one by considering all 125 DNNs in a 5-fold cross validation scheme (see **Methods**).

We evaluated the performance of CCSynergy on two separate datasets namely the Merck dataset^13^ and the one recently published by the Sanger Institute^41^, using two different 5-fold cross validation schemes (CV) (**Fig. 2**). In both CV types, we ensured that any given drug pair in the testing set does not appear in the training set (held-out drug combinations). Furthermore, in the CV2 scheme, we guaranteed that the cell lines in the training set originate from tissues that do not overlap with that of the testing set (held-out tissues). We also investigated cross-data learning to check the potential of CCSynergy to generate experimentally validated predictions. This guided us to generate a database including millions of unexplored (drug pair + cell line) triplets, which we partially validated using its overlap with the DrugComb database^51,52^.

### CCSynergy outperforms state-of-the-art methods on the Merck dataset

We first aimed to evaluate the performance of CCSynergy as compared to its competitors namely DeepSynergy and TranSynergy. Therefore, we applied the CV1 cross validation scheme on the Merck dataset (see **Methods**) and trained the 125 distinct DNNs within the CCSynergy framework. In **Fig. 3a**, we show Pearson correlation coefficient (PCC) between the predicted and real values (averaged among the five folds) for the five CCSynergy methods across the 25 CC spaces. Several important patterns are relevant: *i)* CCSynergy I is clearly outperformed by the other four methods, which implies that gene expression profiles on their own are not strong enough for representing the cell. *ii)* in CCSynergy II-V methods, all 25 CC signatures are highly informative (PCC>0.7), and their relative ranking remains almost the same. For example, E3 always yields the highest PCC, while C2 always stays the lowest. *iii)* CCSynergy II is outcompeted by the other three, but still remains quite close to them. It is remarkable that TF activity on its own could get so close to the signaling pathway-based profiles. *iv)* expression profiles when combined with causal reasoning (CCSynergy III) can yield the same PCC as CRISPR-based essentiality profiling (CCSynergy IV and V). *v)* Integrating the 25 CC spaces by simple averaging always leads to a higher PCC. **Fig. 3b** reveals that all CCSynergy methods (except the first one), when integrating the 25 CC spaces, outcompete both DeepSynergy and TranSynergy. CCSynergy II (0.78) is very close to DeepSynergy (0.77), but CCSynergy III, IV and V yield significantly higher PCC (above 0.81), while TranSynergy (0.69) and CCSynergy I (0.62) clearly lag behind. It is of note that CCSynergy methods cannot significantly surpass DeepSynergy when using each of the 25 CC spaces separately (**Supplementary Fig. S1**) highlighting the fact that it is the integration of the 25 CC spaces, which empowers CCSynergy.

**Figure 3.**
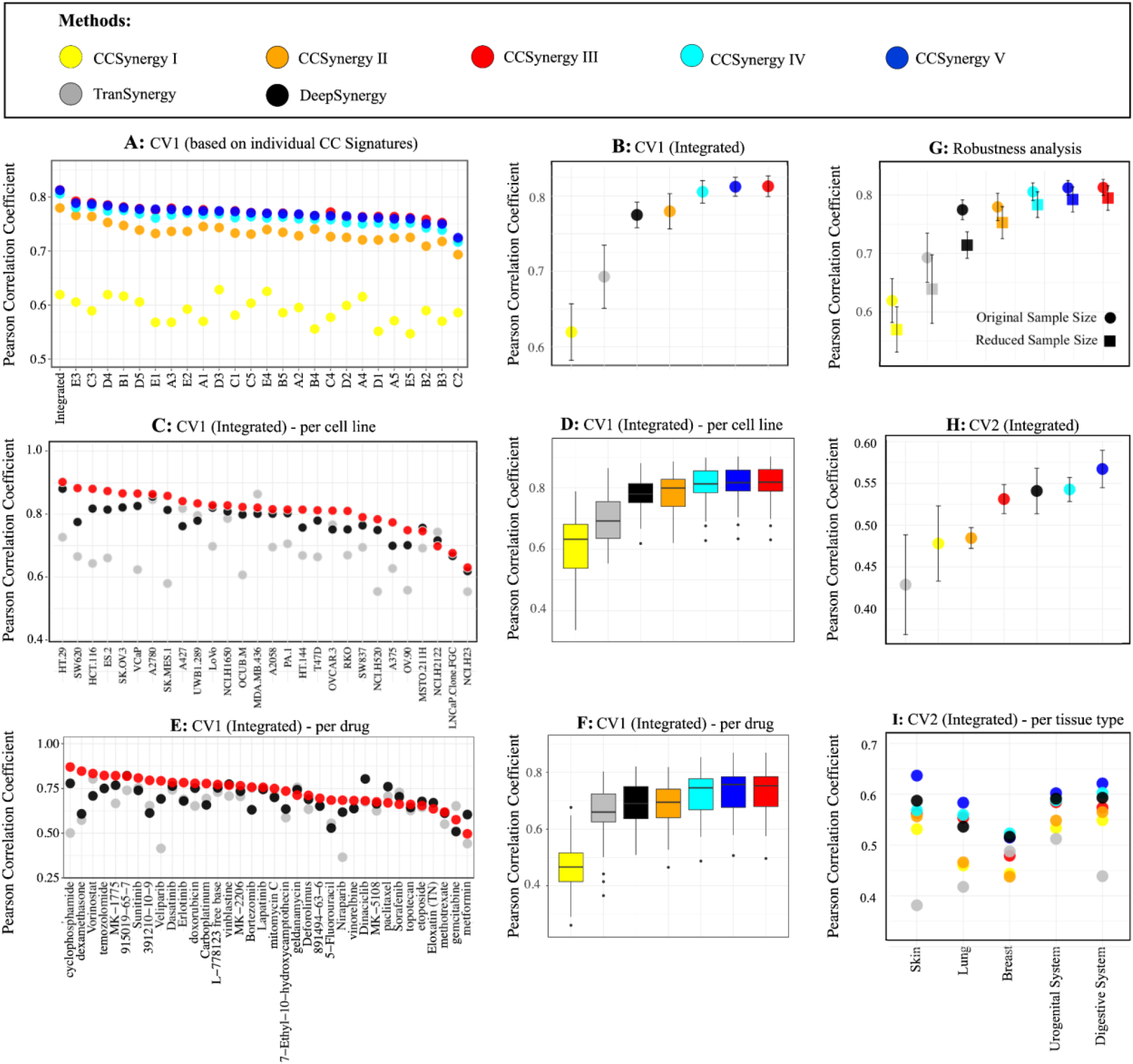
CCSynergy outperforms state-of-the-art drug synergy prediction methods on the Merck dataset. Five versions of CCSynergy method, which differ in their choice of cell line representation methods (See **Fig. 1b** and **Methods**), are color-coded according to the legend (the uppermost box) and are compared against the existing methods such as DeepSynergy^17^ (black) and TranSynergy^18^ (gray). Vertical axes in all panels indicate the Pearson Correlation Coefficient (PCC) between the real and predicted drug synergy values. In panels **A-G**, the PCC scores were calculated within the CV1 scheme, while in panels **H** and **I**, they were measured within the CV2 scheme. In panel **A**, the PCC scores are shown across the 25 CC spaces and also when integrated using simple averaging. In contrast, in panels **B-I** only the integrated PCC scores are shown. Panel **A** only compares the five CCSynergy versions, while in other panels, DeepSynergy and TranSynergy are also included. For clarity purposes, in panels **C** and **E**, only CCSynergy III is shown, while the other four versions are illustrated in **Supplementary Figs. S2** and **S4**, respectively. The circles indicate average PCC across the 5 folds, while the error bars in panels **B, G** and **H** show the corresponding standard deviations. In panel **C**, the PCC scores are calculated for each cell line separately (horizontal axis), and box plots of panel **D** show their distribution (among the cell lines). Similarly, in panel **E**, the PCC scores are shown per drug (horizontal axis), and box plots of panel **F** show their distribution (among the drugs). Moreover, in panel **I**, the average PCC per tissue type (within CV2 scheme) is illustrated. It is important to mention that in panel **G**, the circles indicate the average PCC, when the entire dataset is used (N=14,280), while squares show the average PCC, when a reduced subset of the data is used (N=6,880). Note that all the analyses in this figure are based on the Merck drug synergy dataset (See **Methods**).

We then calculated the PCC scores per cell line to check the consistency of CCSynergy’s performance across different cellular contexts. **Fig. 3c** indicates that in 25 out of 28 cell lines (89.3%), CCSynergy III outcompetes both DeepSynergy and TranSynergy, which is similarly the case for CCSynergy IV and V but not for I and II (**Supplementary Fig. S2**). The distribution of PCC scores across cell lines further confirms the superiority of the CCSynergy III, IV and V when integrating the 25 CC spaces (**Fig. 3d**), but not when using each of them separately (**Supplementary Fig. S3**). We also checked the consistency of CCSynergy performance across different drugs. **Fig. 3e** shows that in 27 out of 36 drugs (75%), the per-drug PCC score of CCSynergy III is above that of DeepSynergy, which is similarly the case for CCSynergy IV and V but not for I and II (**Supplementary Fig. S4**). The distribution of PCC scores across drugs further confirms the superiority of the CCSynergy III, IV and V when integrating the 25 CC spaces (**Fig. 3f**), but not when using each of them separately (**Supplementary Fig. S5**).

Next, we aimed to compare the robustness of these methods to data loss. Thus, we removed 6 drugs and 8 cell lines from the original dataset resulting in a sample of size 6880, which is 48.2% of the original one. We applied the CV1 scheme on the reduced data and calculated the PCC scores for each method. We noted a pronounced reduction of the average PCC scores for DeepSynergy (Δ*PCC=-0*.*061*), TranSynergy (Δ*PCC=-0*.*054*) and CCSynergy I (Δ*PCC=-0*.*049*), while the reductions for CCSynergy II (Δ*PCC=-0*.*027*), III (Δ*PCC=-0*.*018*), IV (Δ*PCC=-0*.*019*) and V (Δ*PCC=-0*.*022*) were significantly less noticeable (**Fig. 3g**). This is especially important and attests to the remarkable robustness and hence more reliable predictions that the integrated CCSynergy framework provides.

Finally, we employed the CV2 scheme to examine the generalizability of these methods for novel cellular contexts. **Fig. 3h** shows the resulting PCC scores for the five CCSynergy methods as compared to the competing ones. The following patterns are germane: *i)* the average PCC scores in CV2 substantially decreased in all methods as compared to the CV1 highlighting the difficulty of drug synergy prediction in novel cellular contexts. *ii)* the PCC score for the CCSynegy II dropped down to the same level as that of the CCSynergy I (0.48) implying that the context-generalizability of the TF-based cell representation is as low as the simple gene-expression based one. *iii)* CCSynergy V (0.57) distinguished itself from the CCSynergy III (0.54) and IV (0.54), which were indistinguishable in the CV1 scheme, and *iv)* CCSynergy V is the only method that significantly outperforms both DeepSynergy (0.54) and TranSynergy (0.43). Furthermore, calculating the PCC scores in the CV2 scheme per tissue type further confirms the superiority of CCSynergy V in all five tissues (**Fig. 3i**). Again, we observed that CCSynergy V has gained its superior performance by integrating the 25 CC spaces as it gets outperformed by DeepSynergy when using each CC signature separately (**Supplementary Fig. S6**), which further highlights the importance of the integrative nature of the CCSynergy framework.

### CCSynergy performs well on the Sanger drug synergy dataset

We then examined the performance of CCSynergy on a new dataset in which the drug synergy is measured differently from the Merck dataset. Thus, we examined the large-scale drug combination screen recently performed in the Sanger institute^41^, which has reported drug synergy in a binary format enabling us to evaluate CCSynergy in a classification setting (See **Methods**). We limited this analysis to CCSynergy III and V, which were the top-performing ones respectively in the CV1 and CV2 schemes on the Merck dataset. Under the CV1 scheme, the corresponding 2×25 DNNs were trained, which output the synergy probability (*θ*) for each testing triplet. This enabled us to calculate the area under the ROC curve (AUC) for these two methods across the 25 CC spaces. **Fig. 4a** shows that *i)* all CC signatures are almost equally informative (AUC ranging between 0.79 and 0.83 in CCSynergy III and between 0.80 and 0.84 in CCSynergy V). *ii)* CCSynergy V yields slightly higher AUC than CCSynergy III across the majority of the CC spaces, and *iii)* integrating the 25 CC spaces (by simple averaging) produces the highest AUC (0.84 in CCSynergy III and 0.86 in CCSynergy V).

**Figure 4.**
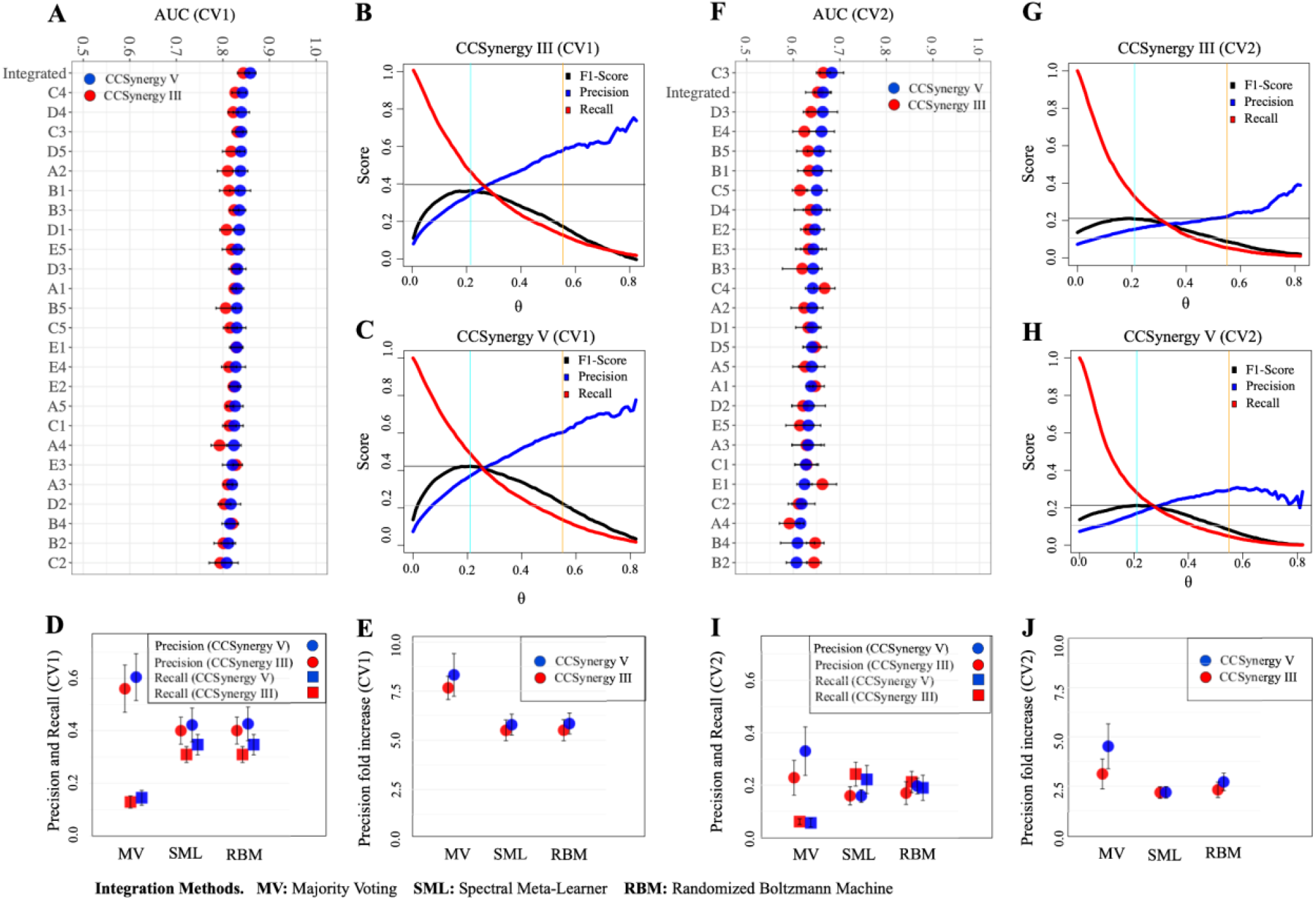
CCSynergy performs well on the Sanger drug synergy dataset. Panels **A-E** show the results obtained under the CV1 scheme, while panels **F-J** show their CV2-based equivalent. **A** and **F)** the average AUC values across the 25 CC spaces plus the integrated one (using simple averaging) are shown as red (CCSynergy III) or blue (CCSynergy V) circles. The curves in panels **B, C, G** and **H** indicate F1-score (black), precision (blue) and recall (red) as a function of the synergy probabilities (*θ*) when using CCSynergy III (CV1: panel **B** and CV2: panel **G**) or CCSynergy V (CV1: panel **C** and CV2: panel **H**). Note that in these four panels, the vertical orange and cyan lines respectively show the *θ*\* and the *θ* maximizing the F1-score. Moreover, the horizontal gray lines respectively show the maximum and half of the maximum F1-score. In panels **D** (CV1), and **I** (CV2), circles and squares indicate respectively the average precision and recall obtained using CCSynergy III (red) or V (blue) after integrating the 25 CC spaces based on the three integration methods mentioned in the horizontal axis, namely: Majority Voting (MV), Spectral Meta-Learner (SML)^53^ and Randomized Boltzmann Machine (RBM)^54^. Similarly, in panels **E** (CV1) and **J** (CV2), the vertical axes indicate the precision fold increase obtained when using CCSynergy III (red) or V (blue) under operation of the three different integration methods. The error bars in all of these panels indicate the standard deviation of the PCC scores across the 5-folds. Note that all the analyses in this figure are based on the Sanger drug synergy dataset (See **Methods**).

Next, we aimed to binarize the outputted synergy probabilities (*θ*) by determining an optimal threshold (*θ*^*^). To this end, we measured F1-score, precision and recall as a function of *θ* for both methods (**Figs. 4b** and **c**). Whereas the recall evidently decreases monotonically by increasing *θ*, we observed that precision increases up to around *θ*=0.6, but then starts to fluctuate. As a common practice in the field, *θ*^*^ is chosen so as to maximize the F1-score, which is a harmonic mean of precision and recall. However, notably precision is much more important than recall for the ultimate goal that drug synergy prediction pursues. Therefore, instead of maximizing F1-score, we selected *θ*^*^ so as to maximize precision subjected to the constraint that F1 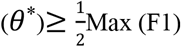. We thus ended up with *θ*^*^=0.55 for both methods. Afterwards, we integrated the binary output of the 25 CC spaces using three different approaches: majority voting (MV), Spectral Meta-Learner (SML)^53^ and Randomized Boltzmann Machines (RBM)^54^. We noted that MV-based integration leads to substantially higher precision and lower recall than SML and RBM methods (**Fig. 4d**). Moreover, **Fig. 4e** indicates more than 7-fold increase of precision (relative to a random classifier) in both CCSynergy methods when using MV and more than 5-fold increase when using SML or RBM. We detected similar patterns, when measuring these metrics per tissue type (**Supplementary Fig. S7**). Importantly, **Supplementary Fig. S8** shows that whereas MV-based integration of the 25 CC spaces always substantially exceeds the single CC-based ones in terms of precision, the SML or RBM based integration always provides superior recall. Furthermore, we observed that all three integration methods always lead to higher convergence between CCSynergy III and V (measured as the Jaccard similarity index; **Supplementary Fig. S9**) than the single-CC based ones. Moreover, intersection of the set of synergistic triplets predicted by CCSynergy III and V culminates in evidently lower recall than either method alone and interestingly higher precision both in the single-CC methods and in the integrated ones (**Supplementary Fig. S8**).

Finally, we performed similar analyses within the CV2 scheme, and first noted that compared to the CV1, the AUC values across all CC spaces have expectedly decreased in both methods, but predictive power to some extent is still preserved (**Fig. 4f**). The average AUC values across the CC spaces varies between 0.60 and 0.68 in CCSynergy V, and similarly between 0.59 and 0.66 in CCSynergy III. We also observed that in both methods, the chosen *θ*^*^=0.55 fulfills the expectations, albeit with a negligible deviation (**Figs. 4g-h**). After discretizing the results, we observed that in all integrative approaches, both precision and recall has decreased in CV2 as compared to the CV1 (**Fig. 4i**). Nevertheless, we can still detect significant enrichment of precision in all integrative scenarios (**Fig. 4j**). SML and RBM based integration yields higher than 2-fold precision increase in both methods, and the VM based one produces even higher enrichment (average 3.1 in CCSynergy III and 4.5 in V). Moreover, the detailed patterns described in CV1 regarding the superiority of the integrative approaches over the single CC-based ones (**Supplementary Figs. S7-S9**), are also similarly detected here (**Supplementary Figs. S10-S12**). Thus, CCSynergy remains helpful, even when applied for predicting drug synergy in novel cellular contexts, and it performs well on an alternative drug synergy dataset, where it is evaluated in a classification setting.

### CCSynergy is of potential to generate experimentally validated predictions

Our next goal was to evaluate a higher-level generalizability of CCSynergy within a cross-dataset learning scheme, which is challenging, especially if drug synergy is measured differently across datasets. This is indeed the case for the Mark and the Sanger datasets, which we used as the training and the testing sets, respectively. This can be regarded as a large-scale experimental validation of the CCSynergy predictions. We ensured that no triplet is shared between the two datasets by considering only the cell lines that were not seen in the Merck data. We distinguished between three scenarios in the Sanger data (**Fig. 5a**) and categorized a given drug combination by checking whether: I) both drugs are seen, II) only one of the drugs is seen, and III) neither drug is seen in the Merck dataset. We then trained 2×25 DNNs, corresponding to 2 methods (CCSynergy III and V) and 25 CC spaces using the entire Merck data as the training set in a classification setting.

**Figure 5.**
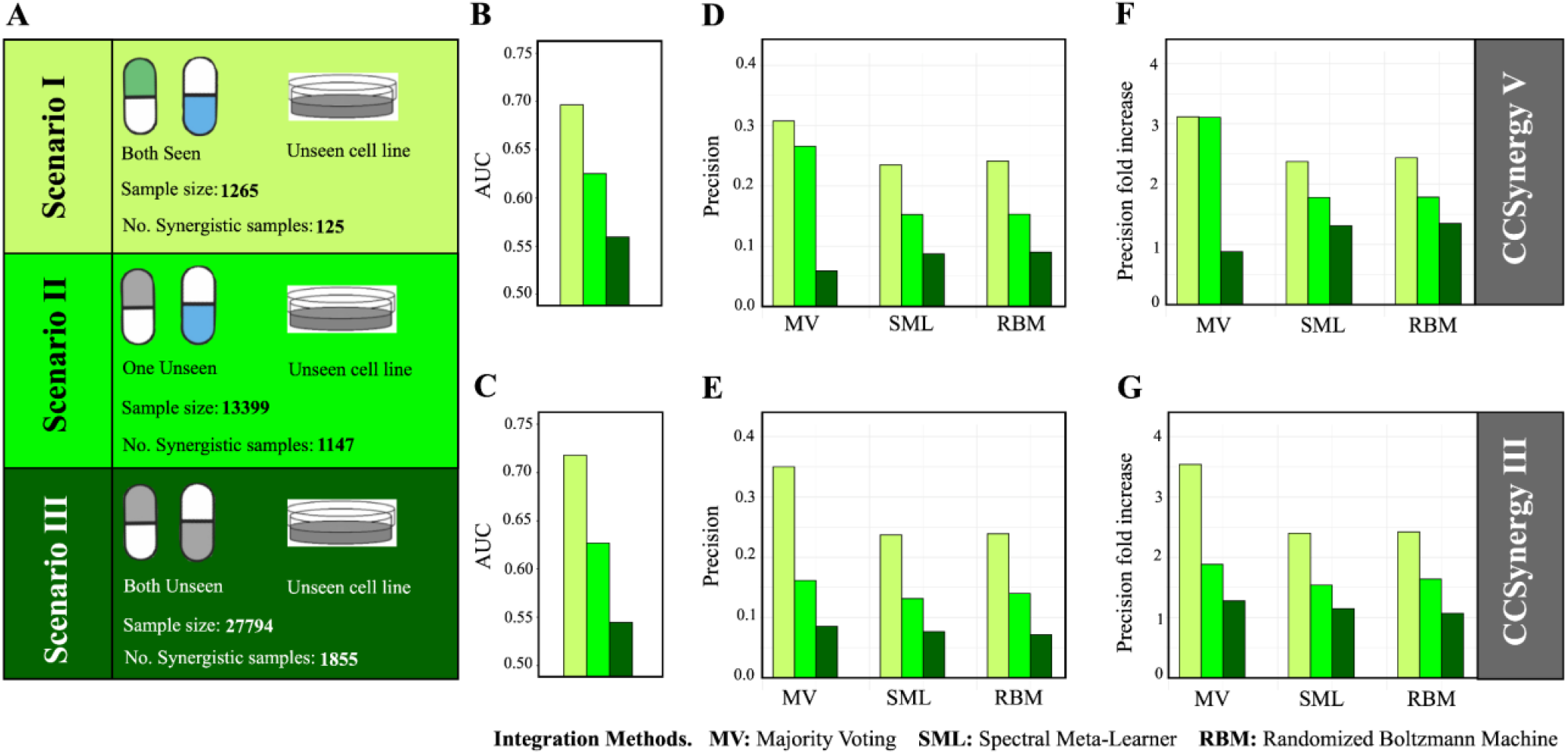
The potential of CCSynergy for generating experimentally validated predictions. In this analysis, we used the Merck dataset as the training and the Sanger dataset as the testing set. Based on their overlap with the Merck data, we have considered three scenarios in the Sanger data and analyzed them separately, which are color-coded and described in panel **A**. The upper (**B, D** and **F**) and lower (**C, E** and **G**) panels were obtained using CCSynergy V and III respectively. The vertical axes indicate the AUC (panels **B** and **C**), precision (panels **D** and **E**), and precision fold increase (panels **F** and **G**). Note that in this analysis, the 25 CC spaces were integrated using simple averaging in panels **B** and **C**, while in the other panels three integrative approaches (horizontal axes in panels **D-G**), namely: Majority Voting (MV), Spectral Meta-Learner (SML)^53^ and Randomized Boltzmann Machine (RBM)^54^ were considered.

After integrating the synergy probabilities (*θ*) of the 25 CC spaces by simple averaging, we calculated the AUC values separately for the above three scenarios. **Figs. 5b-c** show that, in line with our expectations, the AUC values in scenario I were pretty good in both methods (0.70 in CCSynergy V and 0.72 in III). In contrast, in scenario III, they were quite close to the baseline 0.50 for both methods (around 0.55) implying that the model is not much better than a random classifier in cases where neither drug is seen in the training set. However, the good news is that for scenario II, we still detected some predictive power as the AUC values were considerably higher than 0.5 in both methods (0.63).

We then binarized the results and integrated the 25 CC spaces using the three integration approaches (MV, SML and RBM). We detected consistent patterns of precision (**Figs. 5d-e**) and its enrichment (**Figs. 5f-g**) across the three scenarios, regardless of the integration approaches and the CCSynergy methods used. In both scenarios I and II we observed enrichment of precision in all cases, and expectedly the enrichment in the first scenario was always higher than 2-fold and stays consistently above the second one. In the MV-based integration, generally we observed higher enrichment (e.g., more than 3-fold in both scenarios for CCSynergy V) as compared to the SML and RBM, which corroborated our previous observations. However, again we did not see a noticeable departure from the baseline (one-fold) and hence no enrichment of precision for the third scenario (**Figs. 5f-g**). Thus, we conclude that CCSynergy retains a considerable predictive power on unseen cellular contexts in a cross-data learning scheme and hence enhances the potential for generating experimentally validated predictions, provided that at least one of the drugs is seen in the training set.

### CCSynergy generates a compendium of potentially synergistic drug combinations

We showed that CCSynergy is of enhanced potential for generating experimentally validated predictions and so it can facilitate exploration of the untested drug combination space. This motivated us to embark on a voyage to exhaustively explore this space. However, the lack of precision enrichment in the third scenario (**Fig. 5**) cautioned us that CCSynergy could be helpful only in a restricted subspace of drug combinations in which at least one of the drugs is seen in the training set. We thus adjusted our exploration strategy accordingly by focusing on a subspace encompassing the pairing of every single drug (62 drugs: anchor drugs) used in our training data (Sanger dataset) with another pool of drugs that we obtained from the GDSC database^55^ (264 drugs: library drugs). By considering 543 well-characterized cell lines, we ended up with a subspace including 7,786,146 unique triplets that were not tested in the Sanger screen (**Fig. 6a**). We then applied CCSynergy III and V across the 25 CC signatures using the entire Sanger data as the training set in order to predict drug synergy for every triplet in this subspace. After binarizing the outputs using *θ*^*^=0.55, two binary matrices with 7,786,146 rows and 25 columns, were generated. Furthermore, integration of the 25 single-CC results using MV, SML and RBM methods generated three additional binary columns that we added to the final matrices (**Supplementary Tables S1 and S2**). We distinguished between three types of triplets in this space (**Fig. 6b**). Note that although the first two types here are equivalent to the scenarios I and II in the previous analysis, the type III here is not, but rather is an easier to predict version of the type II.

**Figure 6.**
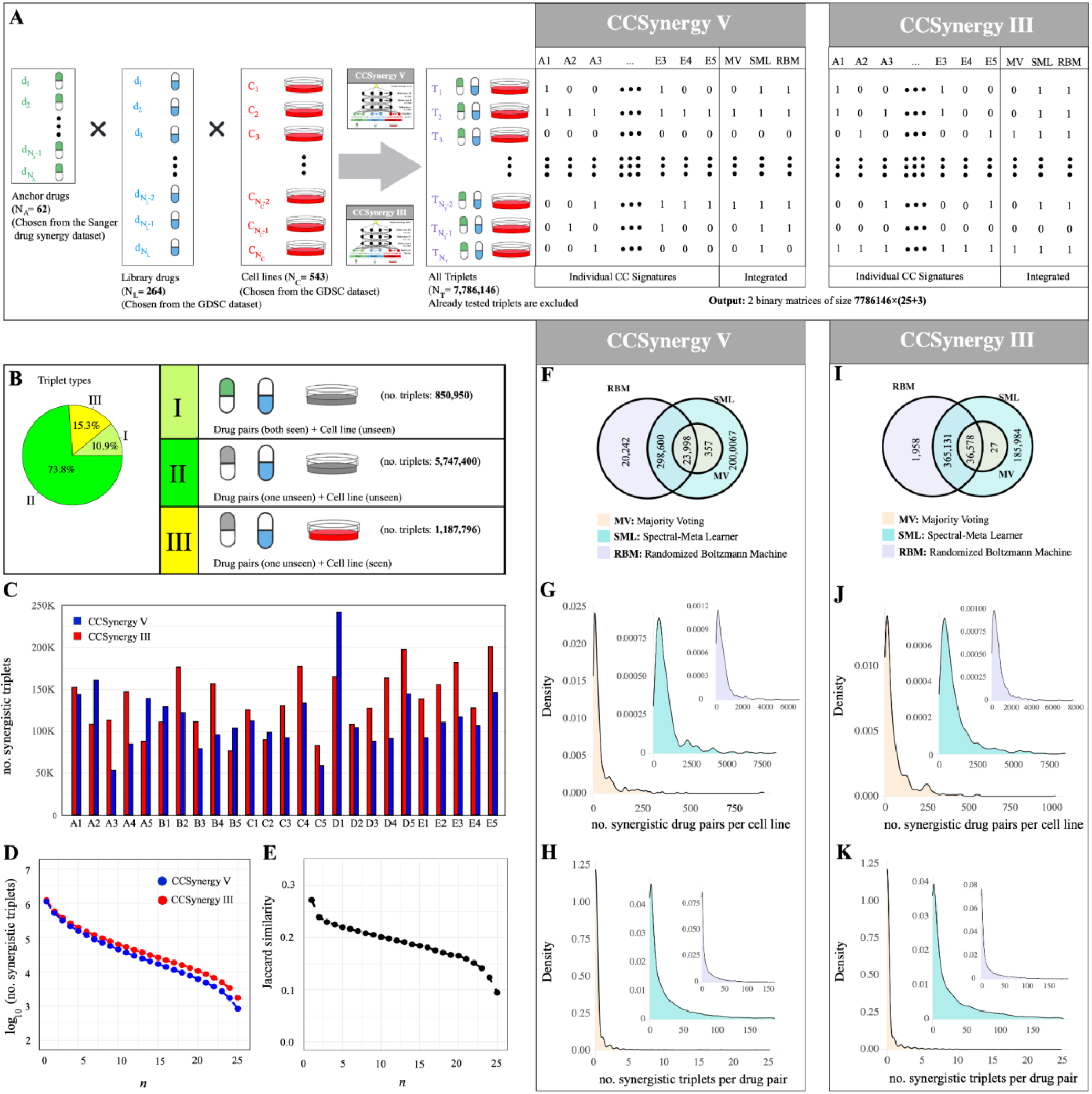
CCSynergy generates a compendium of potentially synergistic drug combinations. **A)** a subspace of the untested drug combination space was considered for exploration, which was constructed by pairing every single drug that was used in the Sanger dataset (62 anchor drugs) with another pool of drugs that were obtained from the GDSC database^55^ (264 library drugs) in 543 well-characterized cancer cell lines. This resulted in a subspace including 7,786,146 unique triplets that were not tested in the Sanger drug combination screen. After training CCSynergy III and V based on the 25 CC signature levels using the entire Sanger dataset as the training set, two binary matrices with 7,786,146 rows and 25 columns were generated. Furthermore, we added three additional columns to these matrices based on the results obtained by MV, SML and RBM integration methods. **B)** We divided the triplets in this subspace into three types based on their overlap with the Sanger dataset. **C)** The bars indicate the number of synergistic triplets across the 25 CC spaces (horizontal axis) identified by CCSynergy III (red) and V (blue). **D)** Each circle indicates the number of synergistic triplets (in logarithmic scale) identified as synergistic in at least *n* CC spaces (horizontal axis) based on CCSynergy III (red) and V (blue). **E)** The Jaccard similarity between the set of synergistic triplets identified by CCSynergy III and the one using CCSynergy V, is shown as a function of *n* (minimum number of CC spaces on which a given triplet is required to be synergistic). **F)** the sets of synergistic triplets after integrating the 25 CC spaces based on MV, SML and RBM using CCSynergy V are identified and the Venn diagram shows the overlap between them. The density plots show the distribution of the number of synergistic triplets identified (using CCSynergy V) **G)** per cell line, and **H)** per drug pair, separately for each of the three integrative approaches. Panels **I, J** and **K** are the equivalent panels based on CCSynergy III.

We enumerated the synergistic triplets for both methods across the 25 CC spaces, which varied between 76,468 (0.98%) and 201,443 (2.59%) in CCSynergy III and between 53,486 (0.68%) and 242,682 (3.11%) in CCSynergy V (**Fig. 6c**). We noted that in both methods a considerable fraction of the triplets is predicted as synergistic at least in one CC space (1,214,794 (15.60%) in CCSynergy III and 1,104,751 (14.18%) in V), but this number declines exponentially by increasing the minimum number (*n*) of required CC spaces (**Fig. 6d**). For example, in CCSynergy V it goes down to 153,091 (1.96%) when *n=5* and to 843 (0.01%) when *n=25*. The agreement between the two methods (CCSynergy III and V), which is measured using the Jaccard similarity of their synergistic triplet sets, in the unseen-cell line scenarios (I and II) is unsurprisingly lower than the seen-cell line type (III) (**Supplementary Fig. S13**). Moreover, the methods diverge further by increasing the minimum number (*n*) of required CC spaces (**Fig. 6e**).

Next, we observed that the MV-based integration of the 25 CC spaces is quite stringent as it identifies only 24,355 synergy cases in CCSynergy V (0.3%), which is a subset of the ones predicted using SML (523,022 (6.7%)) and overlaps strongly (98.5%) also with that of the RBM (342,840 (4.4%)) (**Fig. 6f**). We then checked how synergy is distributed across different cell lines and observed power-law distribution both when using the integration approaches (**Fig. 6g**) or considering the CC spaces alone (**Supplementary Fig. S14**). The implication is that there are few cellular contexts, which are generally more prone to synergy than the others. For example, in the MV-based integration, less than 10% of the cell lines (50 out of 543) account for the majority (12,484 (51.3%)) of the predicted synergies. We observed a similar power law distribution of the number of cell lines providing synergy per drug pair, regardless of the integration methods (**Fig. 6h**) or the single-CC spaces (**Supplementary Fig. S15**) used. This reflects the existence of few drug pairs, which are generally-synergistic independent of the cellular context. For example, the MV-based integration method predicts synergy for Gemcitabine and AZD7762 in 421 out of 543 cell lines (77.5%). Similarly, we found 36 drug combinations (out of 14,483) for which synergy in more than 100 cell lines are predicted. Synergy was found at least in one cell line only for 2,153 drug combinations (15.3%), so for the majority of them (84.7%), synergy was never detected. Analysis of the CCSynergy III results also revealed very similar patterns (**Figs. 6i-k** and **Supplementary Figs. S16-S17**).

To validate (at least partially) our massive predictions, we checked the DrugComb database^51,52^, where the majority of the existing drug combination studies have been amalgamated. We identified 17,472 distinct triplets shared with DrugComb, which includes all three triplet types (**Figs. 7a-b**). The sample size is sufficiently large for a statistical analysis, even though it covers only 0.22% of the triplets in our database. We considered samples with a Loewe score above 9.2 (i.e., the top 10% among the overlapping set) as our reference of true positives. We observed considerable precision enrichment for all three triplet types under RBM (**Fig. 7b**) and SML (**Supplementary Figs. S18a-b**) integration methods. In the MV-based integration scheme, we also observed precision enrichment for triplets of type I, but for types II and III, the sample of MV-based predicted synergies was not of sufficient size (**Supplementary Figs. S18c**). These observations are valid when either CCSynergy III or V method is used, and their intersection leads to even a higher precision enrichment. For example, in the second scenario, the intersection results in 3.10-fold precision enrichment, while the methods alone yield enrichment of 1.35 and 1.87-fold respectively (**Fig. 7b**). **Fig. 7c** lists the 29 triplets that both methods under RBM-integration predict as synergy. As we can see 9 out of the 29 is among the top 10% (true positives), which is considerably larger than 2.9 (expected by chance). Importantly, these true positive cases belong to skin and lung tissues, which were not used in our training set, and so the observed enrichment is not simply an artifact of the choice of tissues in our training set. Furthermore, the enrichment still stays noticeable, if we define synergy more moderately, for example based on the top 25% or 50% triplets. Moreover, we have observed considerable depletion of antagonist ones. We identified only one triplet belonging to the bottom 10%, while the expectation is to observe 2.9 by chance. Thus, in line with our previous observations, the overlap between our database and DrugComb provides additional statistical evidence attesting to the enhanced potential of CCSynergy to generate experimentally validated predictions.

**Figure 7.**
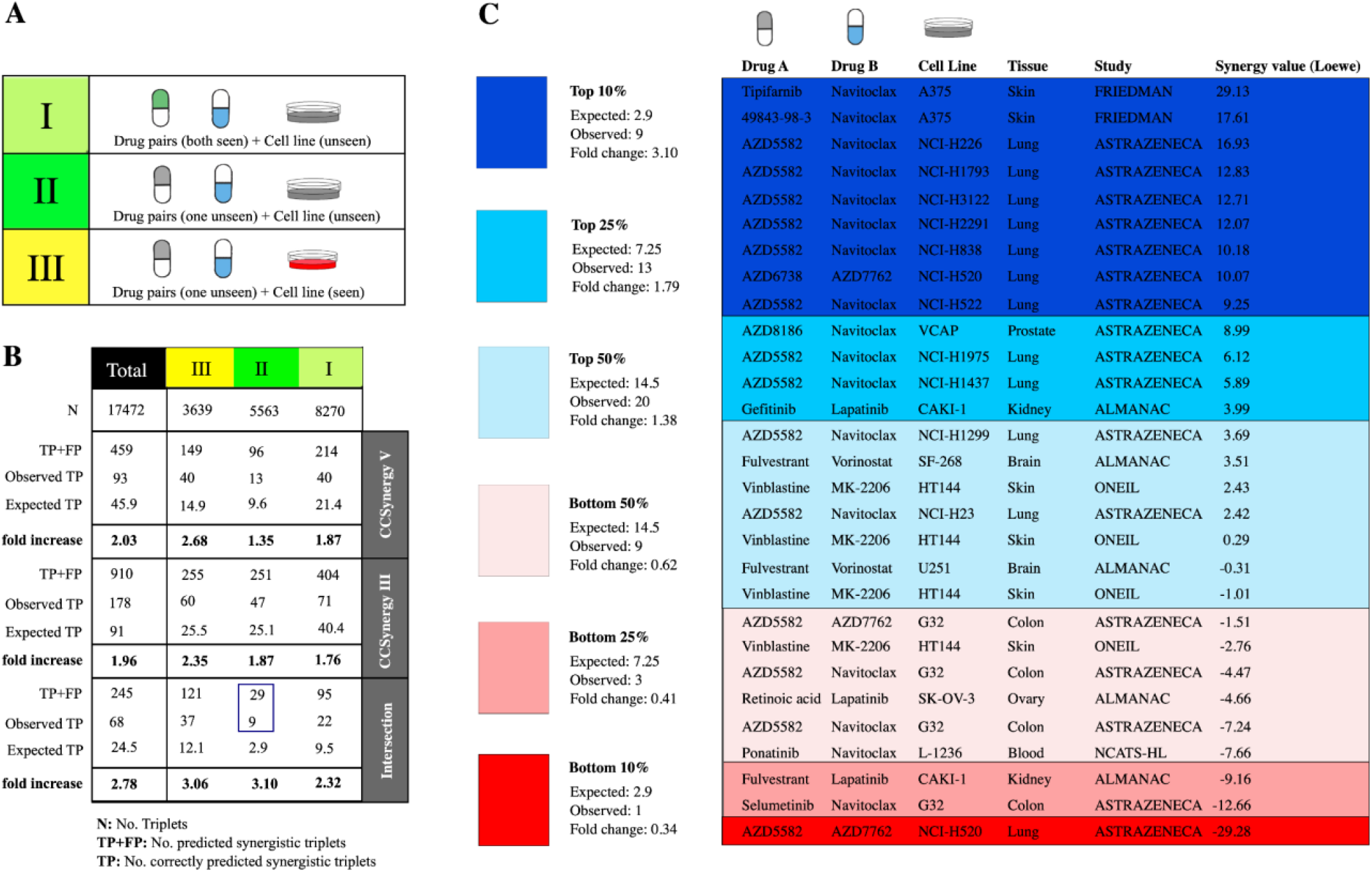
Partial validation of CCSynergy database using its overlap with DrugComb database^51,52^. We identified partial overlap between the triplets considered in the CCSynergy database with those in the DrugComb database. We ranked triplets in this subset in terms of their Loewe synergy score, and considered the top 10% as synergistic (i.e., those with Loewe score > 9.2). **A)** We partitioned the triplets the same as in **Fig. 6b. B)** Number of triplets in the overlapping set (N), number of synergistic triplets predicted by CCSynergy (TP+FP), number of triplets that in both databases are considered as synergistic (i.e., truly synergistic cases: observed TP), the TP that is expected by chance (10% of (TP+FP)), and the precision fold increase, which is basically the ratio of observed by expected TP, are shown for each three scenarios separately and in total (the column names). These measurements have been calculated both for CCSynergy V and III and also their intersection (the row names in the right-hand side). Note that the results in this table were obtained using RBM-based integration of the 25 CC spaces. For the MV or SML-based versions, please see **Supplementary Fig. S18. C)** We have zoomed into the 29 triplets in scenario II identified as synergistic by both CCSynergy III and V methods (i.e., their intersection, which is highlighted by a blue rectangle in panel B). We have listed the drug names, cell lines, tissues, study name and synergy Loewe values for each of the 29 triplets and they are classified and color-coded according to their relative ranking in terms of synergy values in the DrugComb database. For each class, the expected and observed number of true positive cases (triplets identified as synergistic in both DrugComb and CCSynergy databases) along with their corresponding fold changes are specified.

## Discussion

We have introduced CCSynergy, a deep learning framework that we have established to unleash the potential of the Chemical Checker extended bioactivity profiles^40^ for drug synergy prediction. We have proved that the 25 CC spaces provide highly potent representations of drug features and by integrating them, CCSynergy has managed to surpass the state-of-the-art deep learning methods in the field. Moreover, we performed insightful analyses on how to effectively embrace the context-specificity of drug synergy in our predictive models. Firstly, we have demonstrated that down-stream gene expression profiles on their own are not sufficiently informative, but can be substantially upgraded under a causal reasoning framework inferring up-stream signaling-pathway activities (CCSynergy III). Secondly, our analysis revealed that representing cell lines based on genome-wide CRISPR/Cas9 screens, ensures consistent superiority of the model in terms of context-generalizability (CCSynergy V).

Moreover, the fact that CCSynergy performs well on an alternative dataset (Sanger data^41^), where drug synergy was measured differently from the dataset used in the hyper-parameter optimization step (Merck data^13^), confirms its potential for wide applicability. More importantly, we have also demonstrated that compared to its competitors, CCSynergy is remarkably more robust to data loss, which ensures higher reliability and generalizability. Furthermore, we observed considerable precision enrichment when applying CCSynergy in a cross-data learning scheme operated on cell lines that were not seen before. This observation is all the more remarkable, if we consider the fact that drug synergy is notorious for its poor reproducibility across different experimental studies. For example, despite the efforts made in the DrugComb database^51,52^ for standardization and harmonization, the distribution of Loewe scores in the Merck^13^ and NCI-ALMANAC^56^ datasets is still quite different and they are poorly correlated (*PCC=0*.*25*) (**Supplementary Fig. S19**). Thus, the precision enrichments in our cross-data learning analysis deserves to be appreciated and indeed attests to the fact that CCSynergy, to some extent, is of potential for generating experimentally validated predictions and hence can guide future experimental screens by narrowing down the space of untested drug-combinations to a more promising sub-space enriched with true positive cases. This motivated us to cautiously explore this space, which ultimately culminated in a new drug synergy database of unparalleled scale that can be of great assistance for designing follow-up experimental screens.

Nevertheless, as quantified rigorously under the CV2 scheme, our results indicate that there is still ample room for improvement, as current methods are all suboptimal for predicting drug synergy in novel cellular contexts. Importantly, a substantially higher context-generalizability is necessary in order for computational methods to ultimately exert significant clinical impact, especially towards fulfilling the ambitious goals of precision medicine, where the specificity of the cellular contexts plays a decisive role. Thus, the field definitely needs to invest further into exploring innovative strategies on how to represent the cell. However, undoubtedly deep learning methods trained directly on drug combination data, would not be sufficient for this purpose. We strongly believe that insights from single-drug response screens would be the key complement that might also pave the way for detailed mechanistic understanding of drug synergy.

Further directions for methodological improvements exist, both in terms of model architecture and especially with regard to the integration of the 25 CC spaces. For example, instead of CC signatures of type II, the similarity networks inferred from the CC signatures of type I could be employed, which would require using Convolutional Neural Networks (CNNs) instead of DNNs. Alternatively, considering the 25 CC signatures simultaneously in the deep learning framework instead of using them separately is also a valid option. That being said, in the current study, we have already applied three different approaches for integrating the CC spaces, which endow CCSynergy with further flexibility. For example, on the one hand, the majority voting approach leads to a higher precision but lower recall, which can be helpful especially for exploring very large spaces. On the other hand, in scenarios where recall is more important, for example when prioritizing a drug combination list of small or moderate size, applying the SML or RBM methods will be more appropriate.

In summary, we have taken the pioneering step to unlock the potential of Chemical Checker bioactivity profiles by establishing CCSynergy, which is a flexibly integrative framework for context-aware prediction of drug synergy. We anticipate that CCSynergy sparks further methodological developments in the field, inspires new applications, and accelerates exploration of the untested drug combination space.

## Materials and Methods

### Drug synergy data

We analyzed two major drug synergy screens in our study. First, we focused on the Merck dataset^13^, which is based on a large-scale drug combination screen that includes 583 distinct drug pairs, 39 cancer cell lines totaling 22,737 distinct (drug pair + cell line) triplets. We obtained their corresponding Loewe synergy scores from the DrugComb database^51,52^ (https://drugcomb.org/download/; version 1.5), which has accumulated, standardized and harmonized the results of several drug combination screening studies. For a given triplet, we used the average of replicate measurements as the final synergy score. We focused exclusively on drugs, whose CC signatures are available at the Chemical Checker database^40^, and cell lines for which we managed to obtain all the five cell line representations in our study. Therefore, we focused on 36 out of the original 38 drugs and 28 out of the original 39 cell lines totaling 14,280 distinct (drug pair + cell line) triplets (**Supplementary Table 3**).

Secondly, we analyzed the recently published drug combination screen by the Sanger institute (https://gdsc-combinations.depmap.sanger.ac.uk/downloads)^41^, which contains 2,025 pairwise drug combinations and 125 cell lines including breast, colorectal and pancreatic cancer totaling 108,259 distinct triplets. Here, we also focused on a portion of the original dataset, for which both CC signatures and all the cell line representations are available, and so we ended up with 62 drugs, 1177 drug pairs, and 93 cell lines totaling 46,748 distinct triplets (**Supplementary Table 4**). The synergy in the Sanger dataset is reported in a binary format, and the triplets are classified as synergistic (1) or non-synergistic (0) based on the shifts in potency (∆IC50) and in efficacy (∆Emax), which were calculated as the difference between the observed combination response and expected Bliss. The authors have summarized replicate measurements of triplets as synergistic, if half or more of the replicate measurements showed synergy. Moreover, since the design of the Sanger screen distinguishes between the direction of the drug pairs (anchor vs. library), we consider a pair as synergistic if synergy is observed in either direction. Note that in the breast cell lines, the synergy has been measured only in one direction, while for colorectal and pancreatic cancer cell lines, it has been measured in both directions, which largely explains the lower synergy rate that we observed in our analysis for breast (1619 in 30,627 triplets (5.28%)) as compared to those in colorectal (897 in 8349 triplets (10.74%)) and pancreatic (905 in 7772 triplets (11.64%)) cell lines. Note that we employed the Loewe synergy scores of the Merck dataset in a regression setting, while we used binary-scores of the Sanger dataset in a classification setting.

### Drug representation: Chemical Checker extended drug similarity profiles

In order to represent drugs in our deep learning framework, we utilized the extended drug similarity profiles in the Chemical Checker (CC) database (https://chemicalchecker.org/downloads/root)^40^, which systematically catalogs integrated bioactivity data on almost 800,000 small molecules by constructing 25 CC signatures encompassing all levels of bioactivity, which we briefly delineate here (**Fig. 1a**). Level A includes chemical properties of compounds: A1) 2D fingerprints (the 2,048-bit Morgan fingerprints), A2) 3D fingerprints (the 1,024-bit E3FP fingerprints), A3) the Murcko’s scaffold^57^, A4) molecular access system keys representing structural features relevant to medicinal chemistry, and A5) physicochemical parameters of each molecule (e.g. the molecular weight, number of heavy atoms, number of heteroatoms, number of rings, number of aliphatic rings, number of aromatic rings, number of hydrogen bond acceptors, number of hydrogen bond donors and number of rotatable bonds). Level B characterizes the properties of the cellular targets of the compounds: B1) mechanisms of action of approved and experimental drugs, B2) metabolic genes (drug-metabolizing enzymes, transporters and carriers), B3) crystals (protein structures bound to each small molecule), B4) protein binding data, and B5) high-throughput screening bioassays based on bioactivity values from PubChem^58^. Level C covers their pathway/network level properties: C1) small-molecule roles, which have been derived from the ChEBI ontology graph^59^, C2) metabolic pathways, which enable enumerating the set of metabolites in the proximity of a given compound by computing an influence matrix, C3) signaling pathways (the biological pathways that may be affected by the interaction of a molecule with its targets are enlisted), C4) biological processes based on the Gene Ontology Annotation database^60^, and molecules are basically associated with biological process terms based on the annotations of their cellular targets, and C5) interactome, which enables enumerating the set of proteins (on top of the nominal targets) that are influenced by a given compound. Level D pertains to the cellular effects of the drugs: D1) compound-induced gene expression profiles obtained from the L1000 Connectivity Map^28^, D2) small molecule sensitivity profiles of the NCI-60 panel of cancer cell lines^61^, D3) chemical genetics profiles of small molecules screened against ~300 yeast mutants obtained from MOSAIC^62^, D4) morphology profiles obtained from the LINCS data portal, which reports 812 cell image features measured after treatment of cells with ~30,000 compounds^63^, and D5) cell bioassay profiles from ChEMBL^64^. Finally, level E accommodates the clinical effects of the small molecules: E1) therapeutic areas profiled using the ATC classification system codes, E2) Disease indication profiles of drugs obtained from ChEMBL^64^ and RepoDB^65^, E3) side effect profiles of drugs obtained from SIDER^66^, which are expressed as Unified Medical Language System terms, E4) compound-disease association profiles obtained from the Comparative Toxicogenomics Database (CTD)^67^, which contains a medical vocabulary (MEDIC) that is based on the MeSH hierarchy, and E5) drug-drug interaction profiles obtained from DrugBank^68^.

In 22 CC spaces (all except A5, D2 and D4), data are discrete (or discretized) and are expressed as sets of terms such as proteins, pathways, ATC codes and so on. After removing the frequent and rare terms, and down-weighting the less informative ones using frequency–inverse document frequency (TF–IDF) transformation, a dimensionality reduction technique called latent semantic indexing (LSI) has been applied. Similarly, for the continuous data (A5, D2, D4), after robust scaling of the columns, PCA algorithm has been used. This procedure has generated 25 numerical matrices, whose rows correspond to molecules and columns compose the so-called “signatures of type I”, each component of which is orthogonal and sorted by its contribution to the variance of the data. The number of components in each CC space ensures retaining 90% of the variance, and hence the size of CC spaces varies from 500 (A5) to 1500 (D1). Next, to balance the size of the CC spaces, the authors have derived CC signatures of type II, first by building similarity networks from type I signatures, followed by applying a network embedding technique (node2vec^69^). Thus, in the Chemical Checker database^40^, a given drug molecule is ultimately represented by 25 vectors of the same length (128) that we have used as input in our CCSynergy framework (**Fig. 1c**).

### Cell line representation methods

In order to represent cellular contexts in our framework, we examined five different methods as follows (**Fig. 1b**):

**Method I:** This is the simplest approach that we considered as our baseline method, which is simply based on the Affymetrix expression array profiling of the cancer cell lines in the Sanger/MGH GDSC panel (https://www.cancerrxgene.org/gdsc1000/GDSC1000_WebResources/Home.html) that are preprocessed using RMA normalization. The original data includes the basal expression of 19,563 genes for a given cell line, but we first selected the top 1000 genes ranked in terms of their variance across cell lines and then trained an auto-encoder with a single hidden layer including 100 neurons to reduce the dimension from 1000 to 100. The auto-encoder minimizes the mean square error (the difference between predicted and true gene expression vectors) as the loss function. A linear activation function was applied in each layer of the auto-encoder and the weights were initialized randomly and trained using ADAM optimizer with learning rate of 1e-4. The weights were updated after mapping each batch of size 100, and the network training was iterated for 1000 epochs. The final output is available at the **Supplementary Table 5**. Note that the hyper-parameters that we mentioned above were chosen among a combination of choices that we considered in our grid-search towards minimizing the MSE score (**Supplementary Table 6**).

**Methods II and III:** For representing cellular contexts, these two methods rely on the CARNIVAL method^46,70^, which contextualizes signaling networks from gene expression data. Following the standard CARNIVAL pipeline, the RNA-Seq profiles of cancer cell lines (https://cellmodelpassports.sanger.ac.uk/passports) were normalized and compared against their corresponding GTEx normal tissues. Then, the differentially expressed genes were determined using the Limma R package^71^. Next, the transcription factor (TF) activities were estimated using DoRoThea^72^ R package, which systematically catalogs regulons of every transcription factor and outputs the normalized enrichment scores (NES) for TFs based on the transcription status of their corresponding regulons. We did not use the output of DoRoThea directly in our Method II for cell line representation, but rather used it along with the output of PROGENY^73^, which estimates signaling pathway activities based on consensus gene signatures, and the OmniPath-based signed and directed human signaling network^74^, as the inputs of the CARNIVAL method, which infers a signaling subnetwork including up and down-regulated signaling nodes and their interactions. We used the output of the CARNIVAL method in two different ways: whereas **Method II** focuses exclusively on the transcription factors (TFs), **Method III** operates on the level of signaling pathways. In other words, in method II, we focused on the nodes in the CARNIVAL output that are TFs and so we expressed a given cell line as a vector, each element of which belongs to a TF and is +1 (the given TF is an up-regulated node), -1 (the given TF is a down-regulated node) or 0 (the given TF is not present in the inferred network). We chose the top 100 TFs, which are ranked according to their variability across different cell lines (**Supplementary Table 7**). In contrast, for method III, we checked the over-representation of the inferred nodes in 1530 signaling pathways (gene sets) from Reactome^75^ by running gene set analysis using Fisher exact test. We chose 100 signaling pathways (gene sets), which we required i) to be significantly overrepresented (Fisher exact test p-value <0.05) in at least 50 and at most 900 cell lines (out of 1081 cell lines), ii) to be smaller than 100 in size, and iii) to be known as cancer-related. Thus, in method III, a given cell line is represented as a binary vector of length 100, each element of which belongs to a given signaling pathway and is either 1 (significantly overrepresented) or 0 (not overrepresented) (**Supplementary Table 8**).

**Method IV and V:** These two methods both rely on the gene essentiality profiles for representing cell lines (DepMap Public 22Q1: https://depmap.org/portal/download/CRISPR_gene_effect.csv), which contain integrated fold-change depletion values for CRISPR-Cas9 screens performed at either the Sanger or Broad Institutes. The datasets from the two institutes have been batch corrected and combined into integrated datasets that can be analyzed jointly^49^. The dataset covers 908 cell lines and 17,486 genes. We used this dataset in two different ways generating two distinct cell line representation methods: **Method IV**: we first selected 1000 most informative genes, whose depletion fold-change is smaller than (−1) in at least 100 cell lines and at most 800 cell lines. Thus, we initially represented a given cell line using a vector of length 1000 each element of which corresponds to the fold-change depletion of one of the selected 1000 genes. We then reduced the dimension of these vectors from 1000 down to 100 using an auto-encoder with exactly the same architecture and hyper-parameters as in the method I described above (See **Supplementary Table 9**, which includes the final cell line profiles using method IV**). Method V**: we ran an over-representation analysis on the Reactome pathways similarly to what we did in method III (see above) to generate “signaling pathway dependency” profiles for the cancer cell lines. More precisely, for a given cell line, we first identified the set of genes, whose depletion fold-change is smaller than (−1) and then we checked the overrepresentation of this gene set in 1530 signaling pathways (gene sets) from Reactome using Fisher exact test. We chose 100 signaling pathways (gene sets), which we required i) to be significantly overrepresented (Fisher exact test p-value <0.05) in at least 100 and at most 800 cell lines (out of 908), ii) to be smaller than 100 in size, and iii) to be known as cancer-related. Thus, in method III, a given cancer cell line is represented as a binary vector of length 100, each element of which belongs to a given signaling pathway and is either 1 (significantly overrepresented) or 0 (not overrepresented) (**Supplementary Table 10**).

### CCSynergy deep learning framework

Having established our drug and cell line representation methods, we were then able to represent each sample of (drug pair + cell line) triplet by concatenating its corresponding pair of drugs and cell line vectors. It is important to note that we did not concatenate all the 25 different CC signature vectors, neither did we concatenate the 5 different cell line vectors together, but rather generated 125 separate concatenations of the drug pair and cell line vectors of the same size (128+128+100=356), each of which correspond to one of the 125 distinct combinations of the CC spaces and cell line representation methods. Finally, each resulting vector was used as an input in one of the 125 DNNs required to be learnt per a given training set (**Fig. 1c**). Furthermore, it is noteworthy that the synergy scores are order-agnostic and so we needed to represent a given sample twice to account for both directions (i.e., AB and BA). CCSynergy is a feed-forward deep neural network (DNN) and its architecture includes three hidden layers, which propagate information from the input vectors to the output unit, where the predicted synergy score is produced. The number of neurons per layer is *N1=2000, N2=1000* and *N3=500*, which are determined along with other hyper-parameters of the model after considering 256 different choices of hyper-parameter combinations (**Supplementary Table 11**) in our grid-search towards maximizing the average Pearson’s correlation coefficient between the predicted and the real Loewe synergy scores over all 125 distinct DNNs in a drug-combination out 5-fold cross validation setting (CV1: see below) operated on the Merck drug synergy dataset. It is important to highlight the fact that this hyper-parameter optimization step was unavoidably time and resource intensive as we were required to consider all CC spaces and cell line representation methods (256 hyper-parameter choices × 125 DNNs) in order to avoid biasing our analysis towards a specific CC space or a particular cell line representation method. We used linear, ReLU and Tanh activation functions in the first, second and third hidden layers respectively, which are defined as follows:

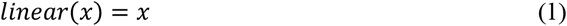

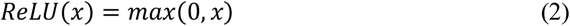

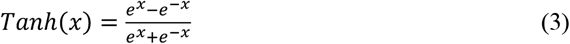

Furthermore, in order to avoid over-fitting, we applied dropout regularization of rate 0.5 and 0.3 respectively in the first and second hidden layers. The weights were initialized based on a truncated normal distribution and trained by minimizing MSE as the loss function according to an Adam optimizer with learning rate 1e-4, *β*_1_ = 0.9 and *β*_2_ = 0.999. We used a batch size of 128 and the maximum number of epochs was set as 1000, but we applied early stopping if the validation loss value did not improve after 10 consecutive epochs. We applied the same DNN architecture and hyper-parameters for analyzing the Sanger drug synergy dataset, but we used binary cross entropy as the loss function, because the synergy values are binary and so the task is defined within a classification setting. Furthermore, we applied a sigmoid activation function (defined below) in the output layer for the classification tasks, while we used the linear activation function in the output layer for the regression ones.

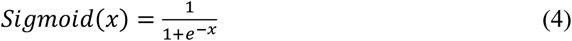

### Cross validation schemes

In our 5-fold cross-validation schemes, we aimed to make sure that a given drug combination in the testing set does not appear in the training set. To do so, we first enumerated all distinct drug pairs and then we randomly divided them into five folds of equal size. We put every (drug pair + cell line) triplets containing a given drug pair in the same fold. The sample of (drug pair + cell line) triplets can be visualized as a matrix each row of which corresponds to a given drug pair and the columns correspond to the cell lines. If we make sure a given drug pair is not repeated in multiple rows (i.e., the rows are unique), a simple row-wise division of the matrix into five folds ensures that the drug pairs in the testing and training sets do not overlap (held-out drug combinations (CV1) (**Fig. 2a**)). In this scenario, five learning cycles are required, in each of which one of the five folds is used as the testing set, and from the remaining four, one is randomly selected as validation set and the other three as the training set. Notably, since our data is obtained from drug combination screens in which every drug pair is tested on the same array of cell lines, the matrix is full and so its row-wise division results in five folds of equal size.

In the second type of 5-fold cross validation (CV2), which is even more strict, we additionally make sure that the tissue of origin corresponding to the cell lines in the testing set do not overlap with that of the training set (held-out tissue). To do so, we needed to divide the matrix not only row-wise but also column-wise, so that all the cell lines in the same testing set belong to the same tissue (**Fig. 2b**). If the cell lines in our data belong to L different tissues, in this CV scheme, 5×L learning cycles are needed. In each cycle, drug pairs of the samples in the testing set must belong to one of the given five folds (the same as in CV1) and also their cell lines must all belong to one of the L tissues (testing tissue). Furthermore, the triplets whose cell lines belong to the given testing tissue must be removed from their corresponding training/validation sets, so that neither rows nor columns are shared between the training and testing sets. Notably, in contrast to the CV1, the size of the testing sets in different cycles of the CV2 scheme may not be equal, because it can vary depending on the number of cell lines belonging to the testing tissue in our samples.

### Method comparison

We compared CCSynergy against the state-of-the-art deep learning methods for anti-cancer drug synergy predictions namely DeepSynergy^17^ and TranSynergy^18^, which have both been applied on the Merck dataset^13^. We needed to re-run both of them, because we have used a subset of the original Merck dataset and also we have obtained the drug synergy scores from the DrugComb database^51,52^, which has standardized and harmonized the original drug synergy scores. We used the same hyper-parameters as what were used in the original papers and we employed exactly the same cross validation schemes as what we used for evaluating the CCSynergy method (CV1 and CV2 described above). DeepSynergy uses connectivity fingerprints, physicochemical and toxicophores for representing drugs, while TranSynergy is based on drug target profiles processed using a random walk with restart (RWR) algorithm. Furthermore, DeepSynergy uses Affymetrix-based gene expression profiles for representing the cell line, while TranSynergy combines gene expression profiles with the gene dependency ones. We used exactly the same drug and cell line features that were used originally, which are pre-computed and downloadable from their code repositories as follows: http://www.bioinf.jku.at/software/DeepSynergy/ (DeepSynergy) and https://github.com/qiaoliuhub/drug_combination (TranSynergy).

### Evaluation metrics

To evaluate the performance of the models in the regression setting, we mainly used Pearson Correlation Coefficient (PCC) between the predicted and the real values. In our 5-fold CV schemes, we obtained PCC of each fold separately and reported their average and standard deviation. In our CV1 scheme, we also reported PCC values per drug and per cell line, in which case we calculated the PCC values using sub-vectors of the original real and predicted vectors, in which only a subset of the data containing the given drug or the cell line is retained. In the CV2 scheme, for a given fold, we merged the results of different tissues and calculated the PCC using the merged vectors. We also measured PCC per tissue, in which case we directly used the predictions made for the given testing set without the need for merging. For performance evaluation in the classification setting, we used the standard area under the ROC curve (AUC), precision, recall and F1-score, which are defined as follows:

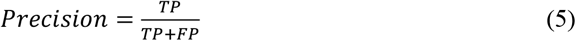

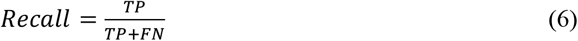

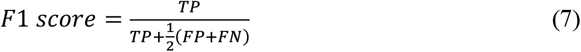

Where: TP, TN, FP and FN are the number of true positive, true negative, false positive and false negative triplets in the given sample. Positive and negative here refers to the triplets predicted as synergistic and non-synergistic respectively. Furthermore, we introduced the so-called “precision fold increase” that is the precision of our predictive model relative to a random classifier, whose precision equals the fraction of positive samples (i.e., synergy rate). Hence, precision fold increase can be formally expressed as:

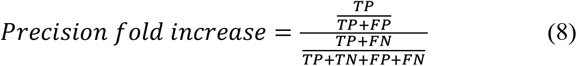

Note that the output of CCSynergy in the classification setting is initially expressed as class probabilities, which are then binarized as synergy (1) or non-synergy (0), if it exceeds our chosen threshold (*θ*^*^=0.55) (**Fig. 4**). We evaluated the performance of CCSynergy both separately for the 25 distinct CC spaces and also integratively. In the regression scheme, we simply used the average of the 25 results to generate the integrated predictions, while in the classification setting we examined three different strategies for integrating the 25 binary results namely: Majority Voting (MV), Spectral Meta-Learner (SML)^53^ and Randomized Boltzmann Machines (RBM)^54^. When comparing different methods, we also used Jaccard similarity index, which is defined as the number of triplets, which both methods consider as synergistic divided by the number of triplets, which at least one of the methods considers as synergistic.

## Statistical analyses

All statistical analyses were performed using the R software (v4.1.2 https://www.r-project.org/).

## Code availability

CCSynergy codes and data will be available upon publication.

## Acknowledgement

This work was supported by funding from the National Institute of Health (grants numbers: NIH R01GM123037, U01AR069395, R01CA241930, and NSF 2217515). The authors would like to thank Julio Saez-Rodriguez and Rosa Hernansaiz-Ballesteros for sharing their CARNIVAL data. Furthermore, S-R.H appreciates inspiring discussions with Mathew Garnett regarding the drug combination resource that has recently been published by his team. SRH is also thankful to Mahya Mehrmohamadi for helpful discussions and Narjes Rohani for technical assistance. Computational resources for this project were provided by Texas Advanced Computing Center (TACC). Funding for open access charge is provided by Dr. & Mrs. Carl V. Vartian Chair Professorship Funds to Dr. Zhou from the University of Texas Health Science Center at Houston.

## Author contributions

S-R.H. and X.Z. conceived the project. S-R.H. designed and performed the analyses and wrote the manuscript. X.Z. acquired funding. Both authors discussed the results, read and approved the final manuscript.

