## Supplementary for "Predicting anti-cancer drug synergy using extended drug similarity profiles"

### Supplementary Tables:

**Supplementary Table S1: CCSynergy database (part I).** This is the CCSynergy database generated using the CCSyngy V method. Note that due to space constraints, we have retained 1,822,681 triplets (out of the 7,786,146), which are predicted to be synergistic at least in one of the CC spaces by either CCSynergy V or III.

**Supplementary Table S2: CCSynergy database (part II).** This is the CCSynergy database generated using the CCSyngy III method. Note that due to space constraints, we have retained 1,822,681 triplets (out of the 7,786,146), which are predicted to be synergistic at least in one of the CC spaces by either CCSynergy V or III.

**Supplementary Table S3: Merck drug synergy dataset.**

**Supplementary Table S4: Sanger drug synergy dataset.**

**Supplementary Table S5: CCSynergy I cell line representation matrix** (each column belongs to a cell line)**.**

**Supplementary Table S6: Hyper-parameter optimization for the auto-encoder.** (See the next page)

**Supplementary Table S7: CCSynergy II cell line representation matrix** (each column belongs to a cell line and each row corresponds to one of the 100 transcription factors)**.**

**Supplementary Table S8: CCSynergy III cell line representation matrix** (each column belongs to a cell line and each row corresponds to one of the 100 signaling pathways)**.**

**Supplementary Table S9: CCSynergy IV cell line representation matrix** (each column belongs to a cell line)**.**

**Supplementary Table S10: CCSynergy V cell line representation matrix** (each column belongs to a cell line and each row corresponds to one of the 100 signaling pathways)**.**

**Supplementary Table S11: Hyper-parameter optimization for the DNN architecture.** (See the next page)

*Note that supplementary tables will be available upon publication.

### Supplementary Figures:

**Supplementary Figures S1-S19:** See pages 3-18.

| Parameter | Options considered | Chosen option |
|---|---|---|
| Optimizer | [ADAM, SGD] | ADAM |
| Activation function | [Linear, ReLU, Sigmoid] | Linear |
| Learning rate | [0.001, 0.0001, 0.00001] | 0.0001 |
| Batch size | [50, 100, 200, 300] | 100 |

**Supplementary Table S6: Hyper-parameter optimization for the auto-encoder.**

| Parameter | Options considered | Chosen option |
|---|---|---|
| N1 | [500,1000,2000,3000] | 2000 |
| N2 | [500,1000,2000,3000] | 1000 |
| N3 | [50,100,300,500] | 500 |
| Learning rate | [0.001,0.0001] | 0.0001 |
| Batch size | [128,512] | 128 |

**Supplementary Table S11: Hyper-parameter optimization for the DNN architecture.**

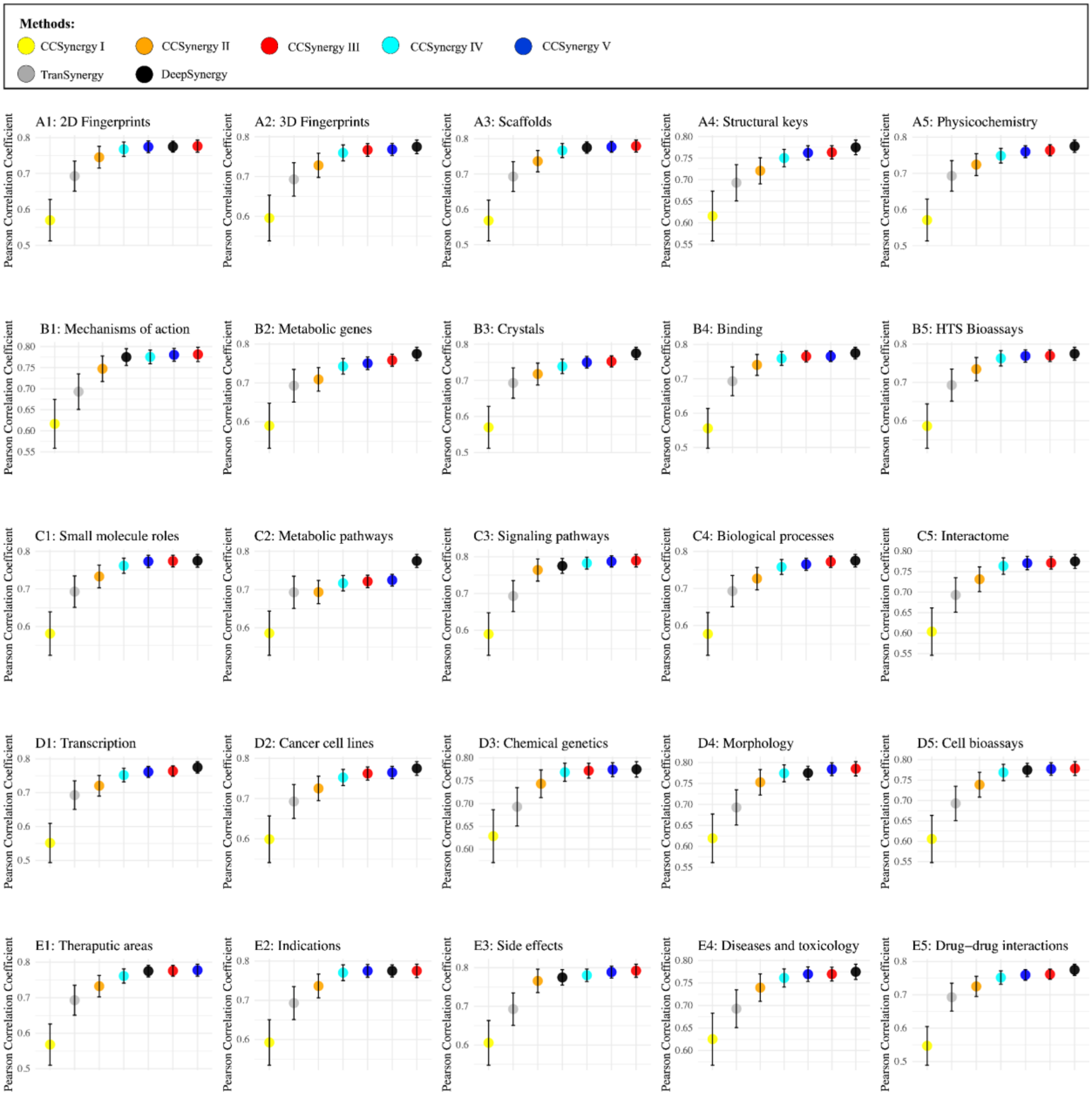


**Supplementary Figure S1. CCSynergy cannot outcompete DeepSynergy when using CC spaces separately (CV1 scheme).** Each panel corresponds to one of the 25 CC spaces and the circles and error bars respectively indicate the average and standard deviation of the PCC scores among the 5-folds within CV1 scheme. Each colorful circle belongs to one of the five CCSynergy methods, which are color-coded according to the legend (uppermost panel), and are compared against DeepSynergy (black) and TranSynergy (gray). This analysis was performed on the Merck dataset.

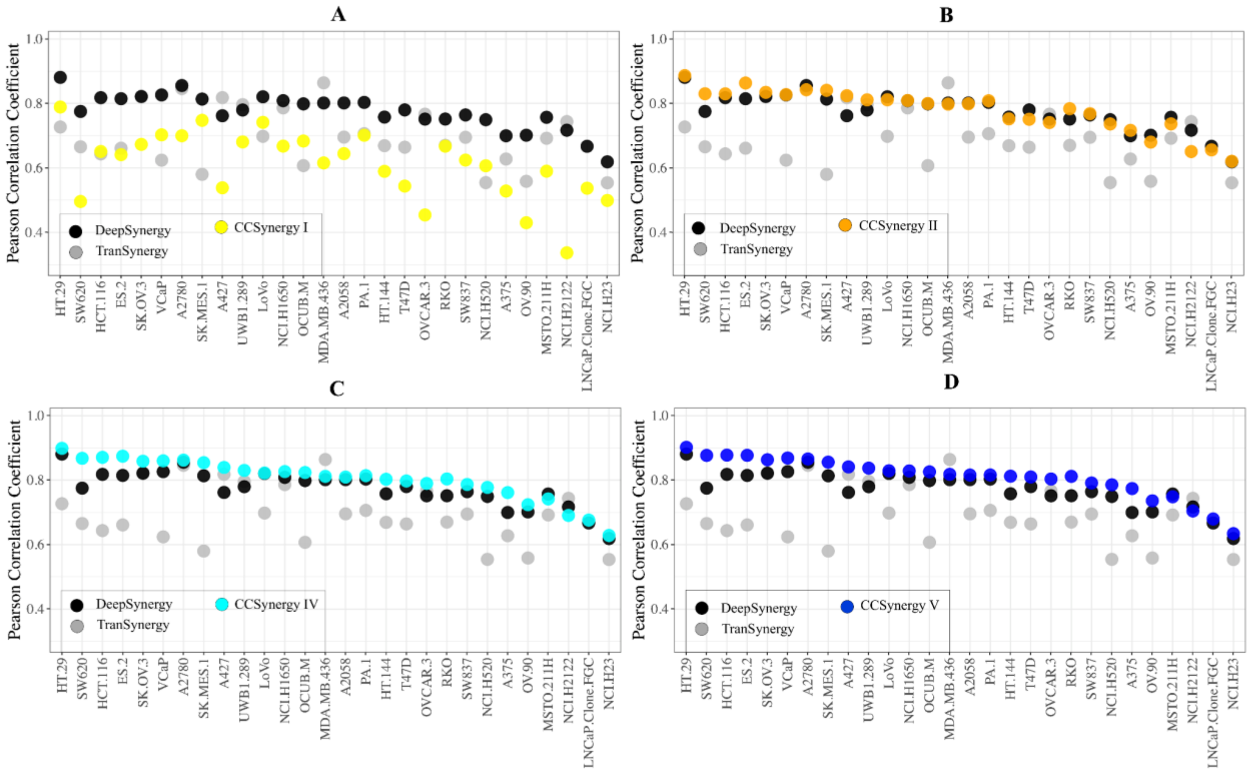


**Supplementary Figure S2. Consistency of the CCSynergy performance across different cell lines.** In each panel, one of the CCSynergy methods is compared against DeepSynergy and TranSynergy. Circles indicate the average PCC score (among the 5-folds) across different cell lines (horizontal axis), which are color-coded according to the legends. This analysis was performed on the Merck dataset.

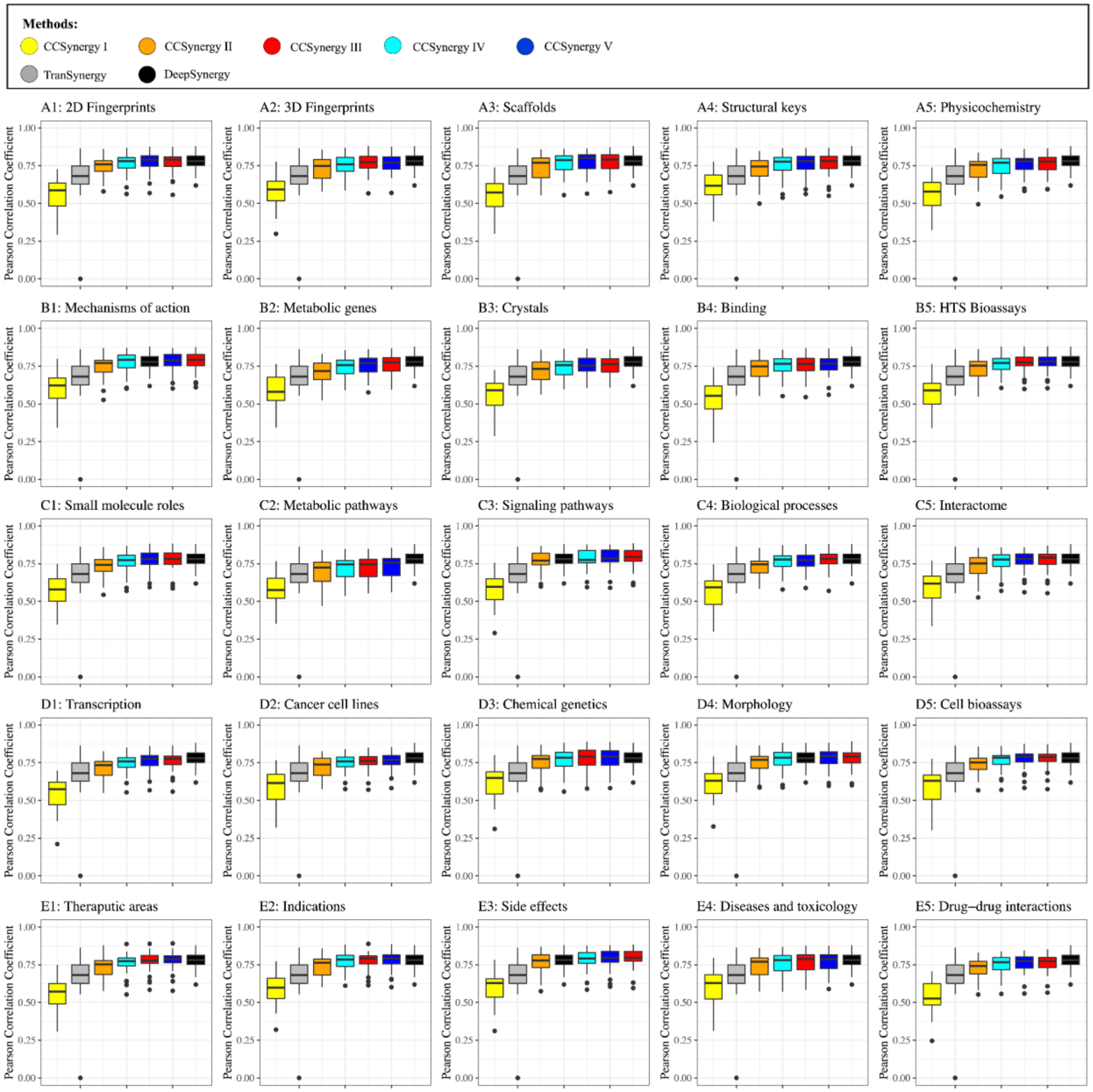


**Supplementary Figure S3. CCSynergy cannot outcompete DeepSynergy when using CC spaces separately (per-cell line analysis).** Each panel corresponds to one of the 25 CC spaces and the box plots show the distribution of the PCC scores across different cell lines. Each colorful box belongs to one of the five CCSynergy methods, which are color-coded according to the legend (uppermost panel), and are compared against DeepSynergy (black) and TranSynergy (gray). This analysis was performed on the Merck dataset.

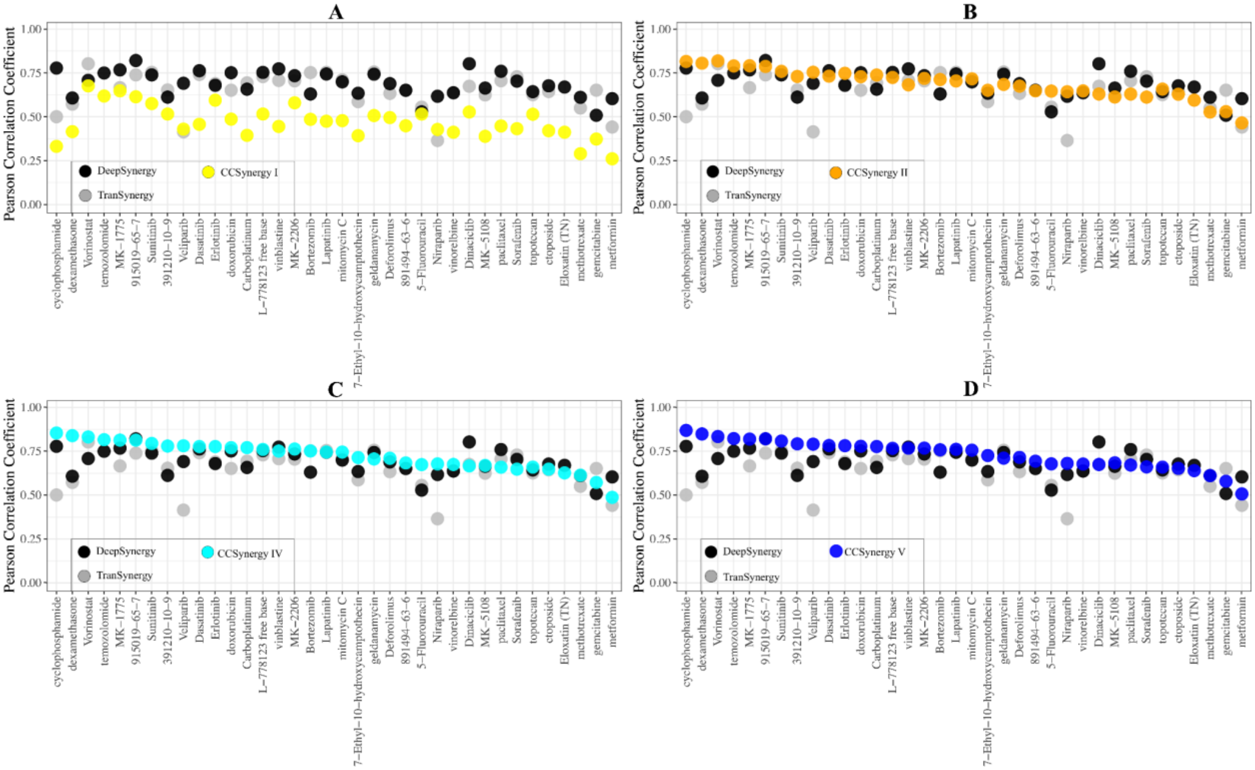


**Supplementary Figure S4. Consistency of the CCSynergy performance across different drugs.** In each panel, one of the CCSynergy methods is compared against DeepSynergy and TranSynergy. Circles indicate the average PCC score (among the 5-folds) across different drugs (horizontal axis), which are color-coded according to the legends. This analysis was performed on the Merck dataset.

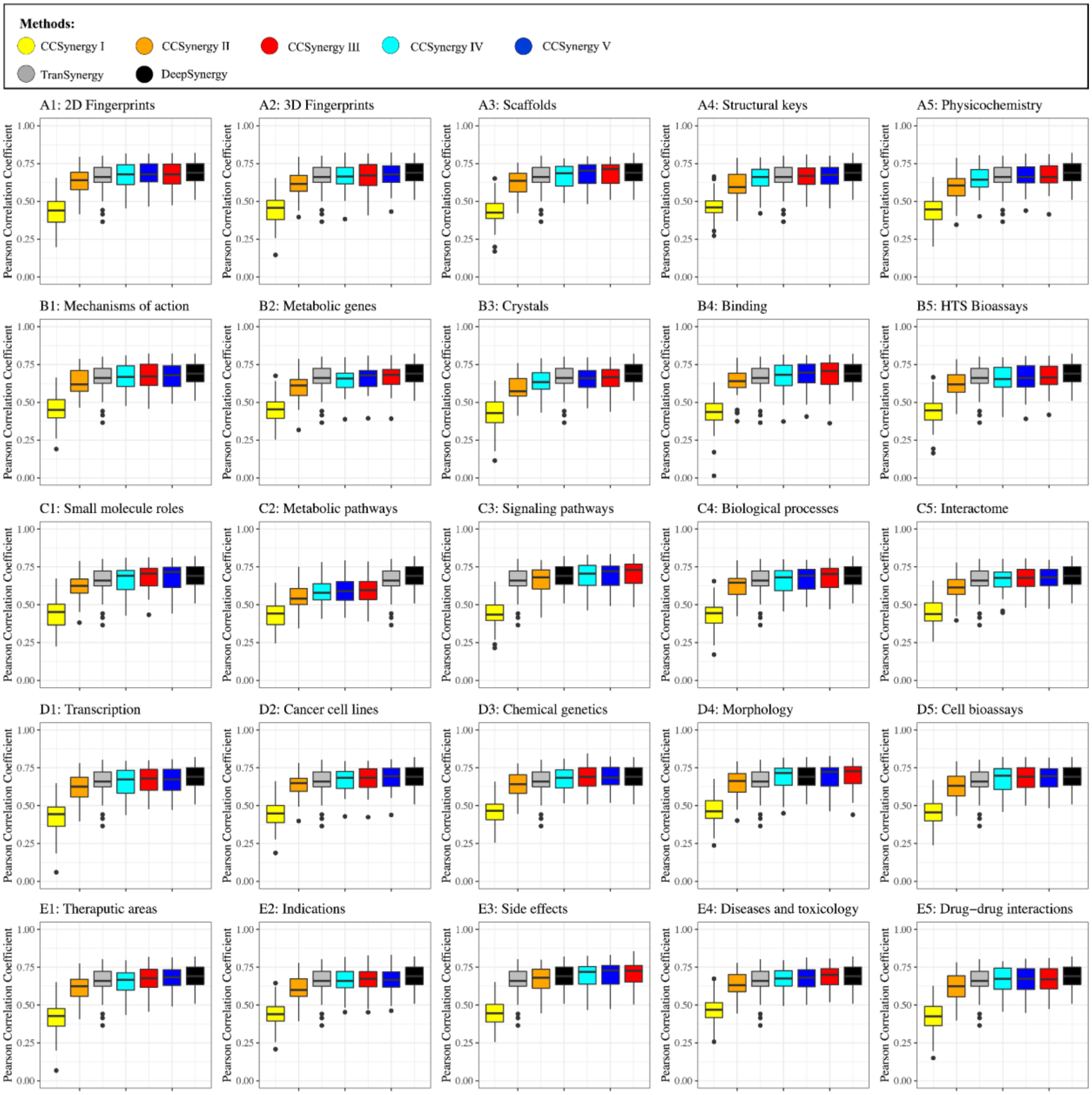


**Supplementary Figure S5. CCSynergy cannot outcompete DeepSynergy when using CC spaces separately (per-drug analysis).** Each panel corresponds to one of the 25 CC spaces and the box plots show the distribution of the PCC scores across different drugs. Each colorful box belongs to one of the five CCSynergy methods, which are color-coded according to the legend (uppermost panel), and are compared against DeepSynergy (black) and TranSynergy (gray). This analysis was performed on the Merck dataset.

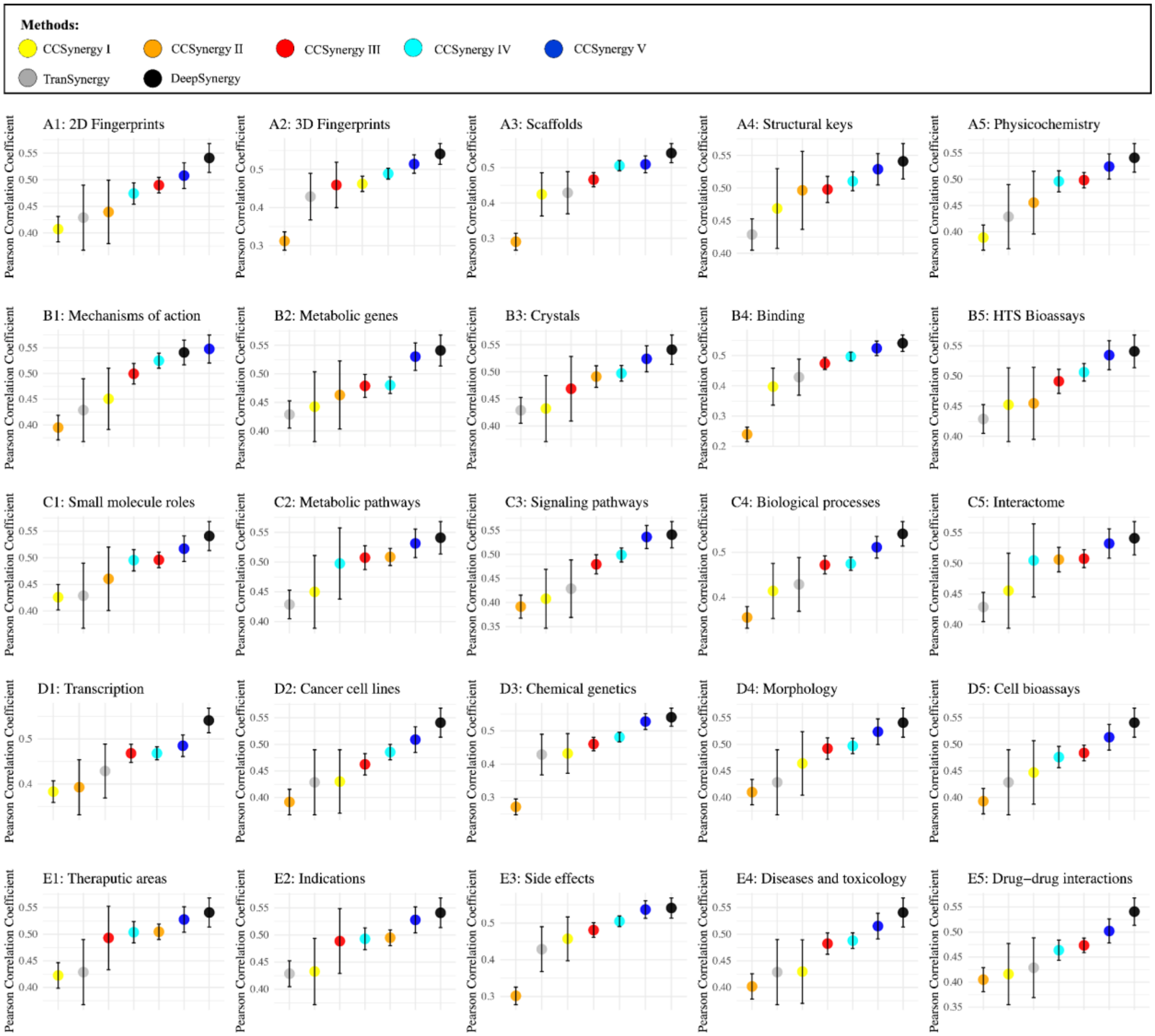


**Supplementary Figure S6. CCSynergy cannot outcompete DeepSynergy when using CC spaces separately (CV2 scheme).** Each panel corresponds to one of the 25 CC spaces and the circles and error bars respectively indicate the average and standard deviation of the PCC scores among the 5-folds within CV2 scheme. Each colorful circle belongs to one of the five CCSynergy methods, which are color-coded according to the legend (uppermost panel), and are compared against DeepSynergy (black) and TranSynergy (gray). This analysis was performed on the Merck dataset.

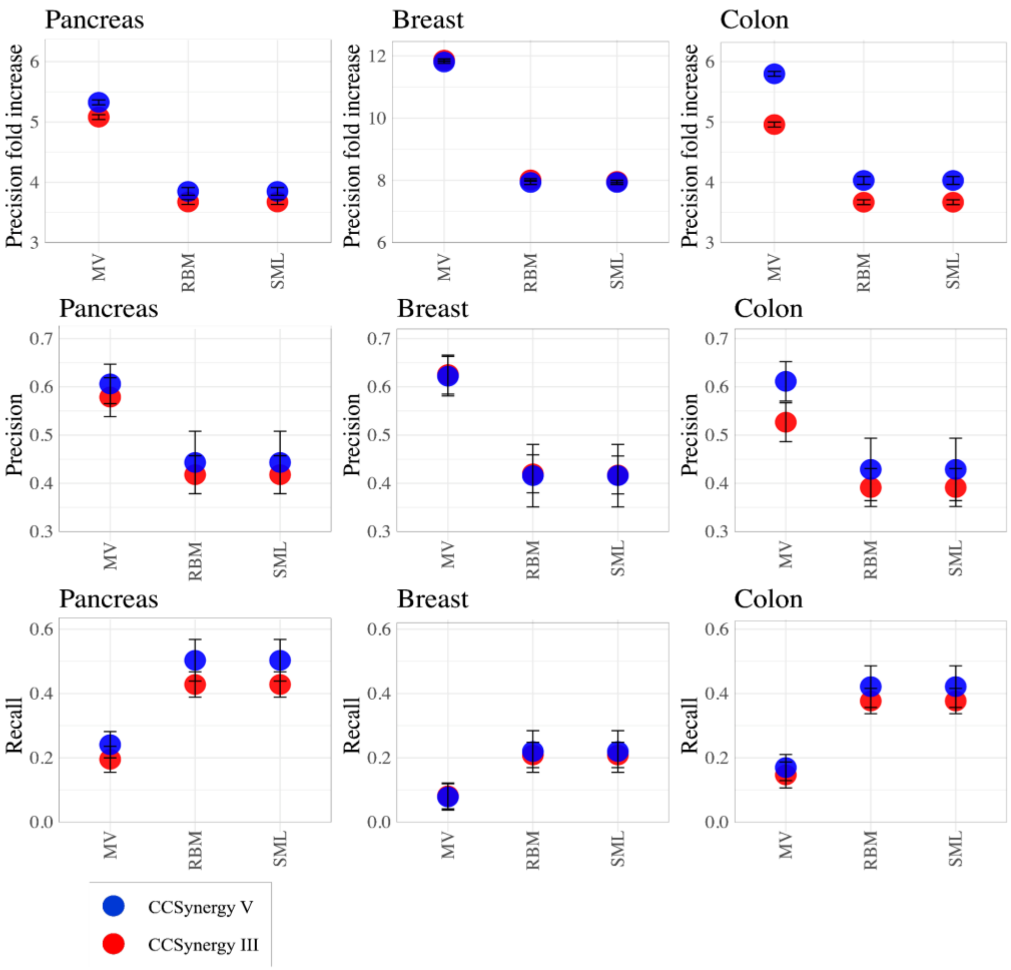


**Supplementary Figure S7. Performance of CCSynergy (integrated) methods (CV1 scheme) across tissue types.** Vertical axes indicate precision fold increase (upper panels), precision (middle panels) and recall (lower panels). The circles and error bars respectively show the average and standard deviation of the metrics (vertical axes) among the 5-folds within the CV1 scheme using CCSynergy III (red) or V (blue), calculated separately for pancreas (left), breast (middle) and colon (right) tissues. The horizontal axes indicate the methods used for integrating the 25 CC spaces, namely Majority Voting (MV), Spectral Meta-Learner (SML) and Randomized Boltzmann Machines (RBM). This analysis was performed on the Sanger dataset.

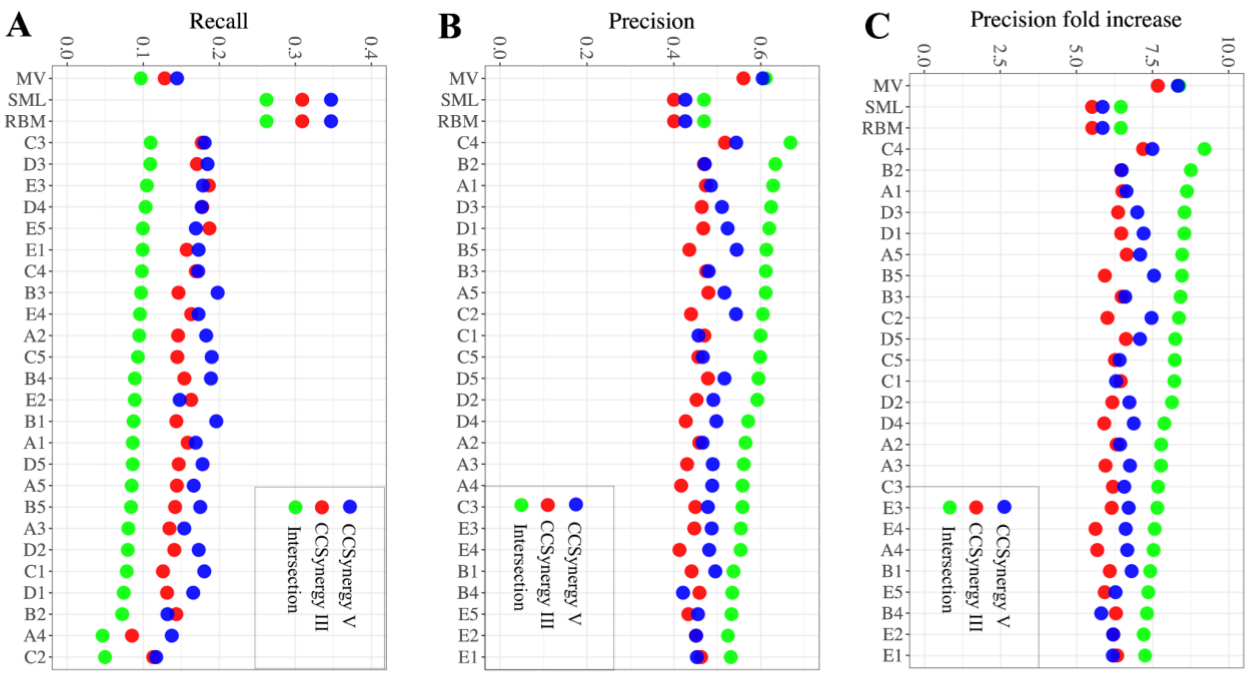


**Supplementary Figure S8. Performance of CCSynergy methods (CV1 scheme) evaluated separately across the 25 CC spaces versus the integrated ones.** Vertical axes indicate A) recall, B) precision, and C) precision fold increase averaged among the 5-folds within the CV1 scheme using CCSynergy III (red) and V (blue) or their intersection (green), calculated separately across the 25 CC spaces and also integratively based on Majority Voting (MV), Spectral Meta-Learner (SML) and Randomized Boltzmann Machines (RBM). This analysis was performed on the Sanger dataset.

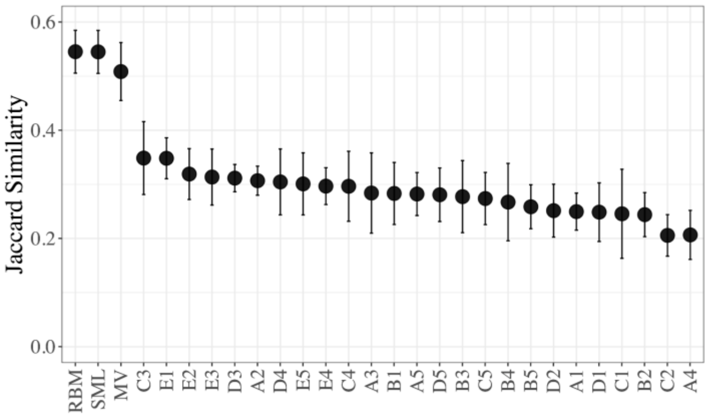


**Supplementary Figure S9. Agreement between CCSynergy III and V (CV1).** Vertical axis indicates the Jaccard similarity (JS) between the set of synergistic triplets predicted by CCSynergy III with that of CCSynergy V. Circles and error bars respectively indicate the average and standard deviation of the JS value among the 5-folds within the CV1 scheme across each of the 25 CC spaces (horizontal axis) and their integration based on Majority Voting (MV), Spectral Meta-Learner (SML) and Randomized Boltzmann Machines (RBM). This analysis was performed on the Sanger dataset.

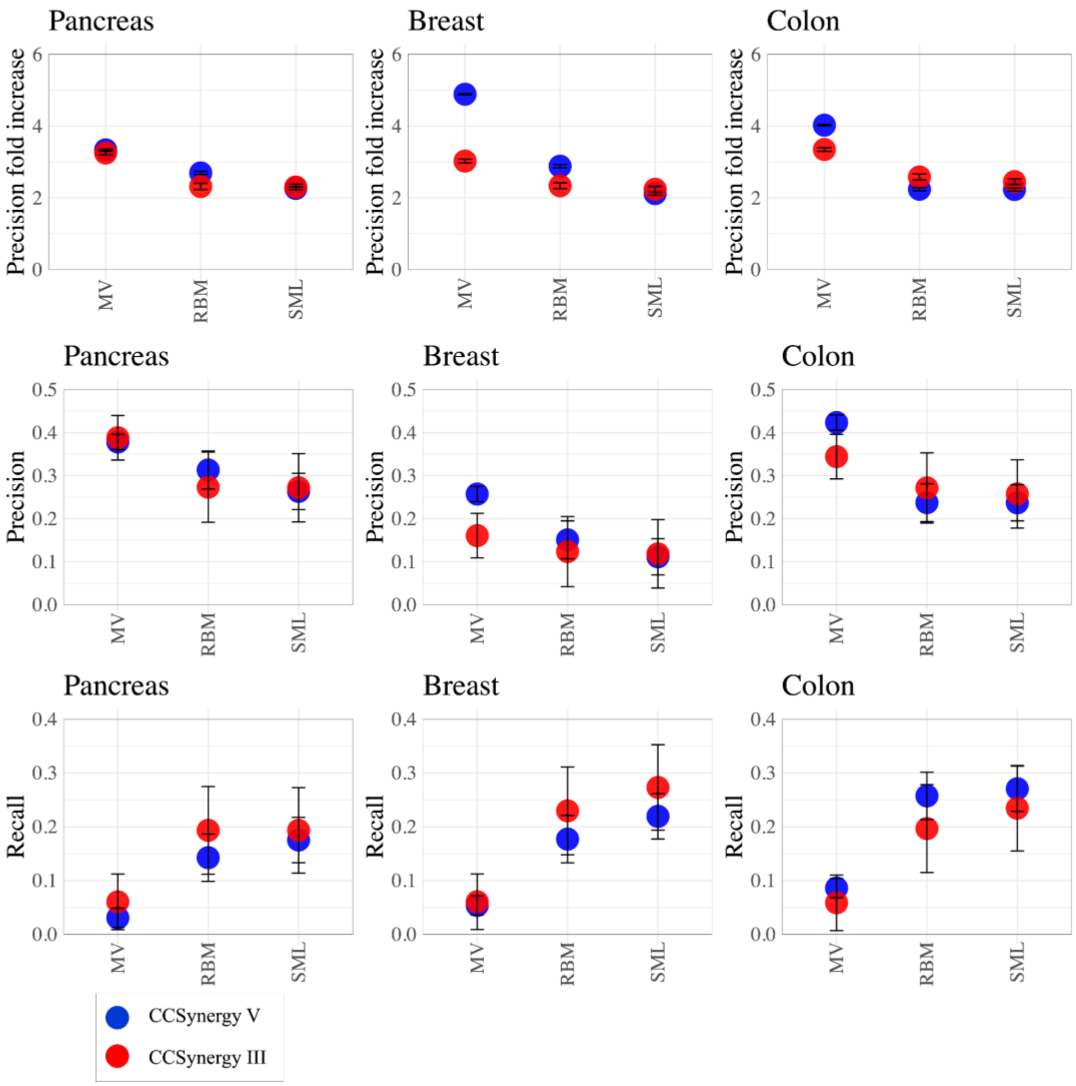


**Supplementary Figure S10. Performance of CCSynergy (integrated) methods (CV2 scheme) across tissue types.** Vertical axes indicate precision fold increase (upper panels), precision (middle panels) and recall (lower panels). The circles and error bars respectively show the average and standard deviation of the metrics (vertical axes) among the 5-folds within the CV2 scheme using CCSynergy III (red) or V (blue), calculated separately for pancreas (left), breast (middle) and colon (right) tissues. The horizontal axes indicate the methods used for integrating the 25 CC spaces, namely Majority Voting (MV), Spectral Meta-Learner (SML) and Randomized Boltzmann Machines (RBM). This analysis was performed on the Sanger dataset.

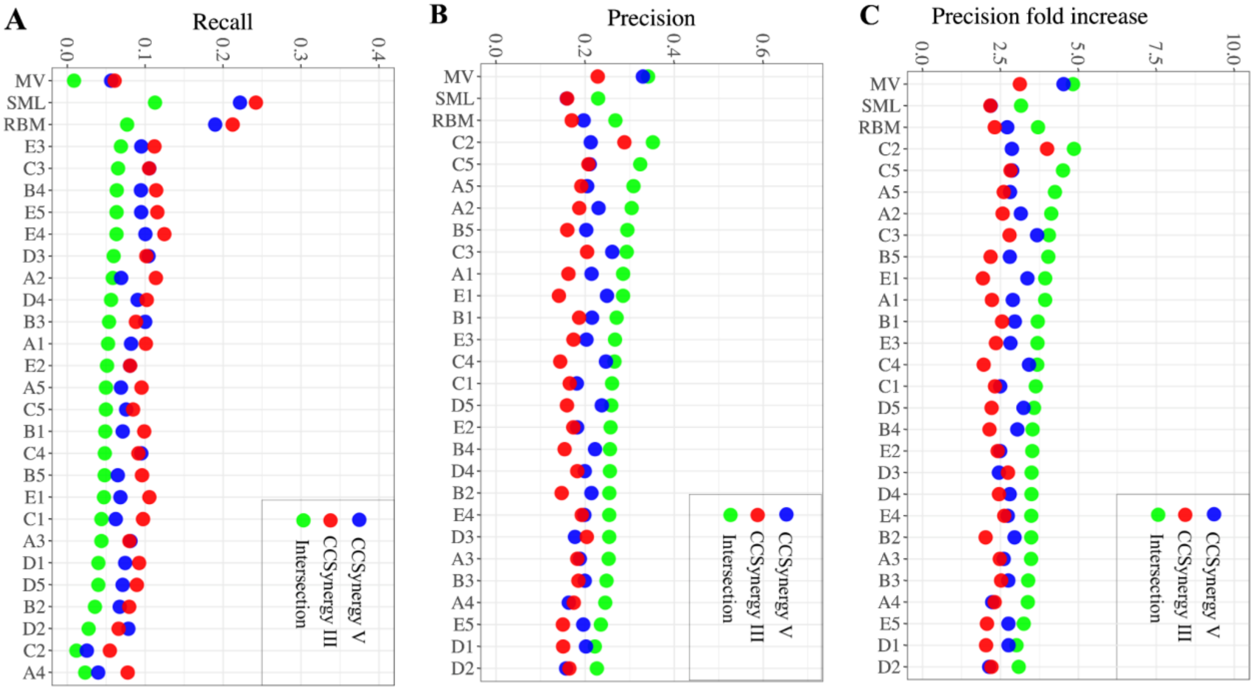


**Supplementary Figure S11. Performance of CCSynergy methods (CV2 scheme) evaluated separately across the 25 CC spaces versus the integrated ones.** Vertical axes indicate A) recall, B) precision, and C) precision fold increase averaged among the 5-folds within the CV2 scheme using CCSynergy III (red) and V (blue) or their intersection (green), calculated separately across the 25 CC spaces and also integratively based on Majority Voting (MV), Spectral Meta-Learner (SML) and Randomized Boltzmann Machines (RBM). This analysis was performed on the Sanger dataset.

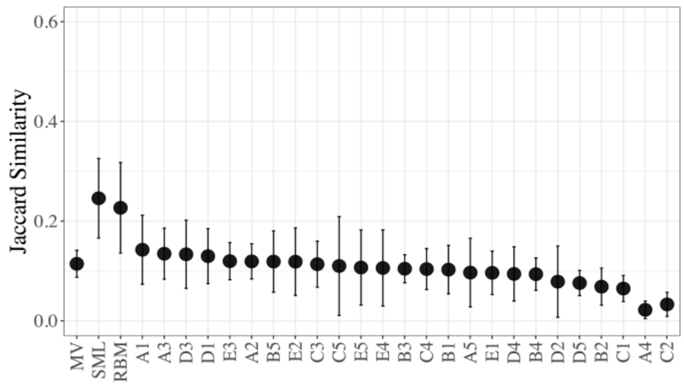


**Supplementary Figure S12. Agreement between CCSynergy III and V (CV2).** Vertical axis indicates the Jaccard similarity (JS) between the set of synergistic triplets predicted by CCSynergy III with that of CCSynergy V. Circles and error bars respectively indicate the average and standard deviation of the JS value among the 5-folds within the CV2 scheme across each of the 25 CC spaces (horizontal axis) and their integration based on Majority Voting (MV), Spectral Meta-Learner (SML) and Randomized Boltzmann Machines (RBM). This analysis was performed on the Sanger dataset. Note that MV-based JS value is not better than the single-CC based methods here unlike in CV1, which might simply be a consequence of the small sample size due to the very low recall (**Supplementary Fig. S11A**).

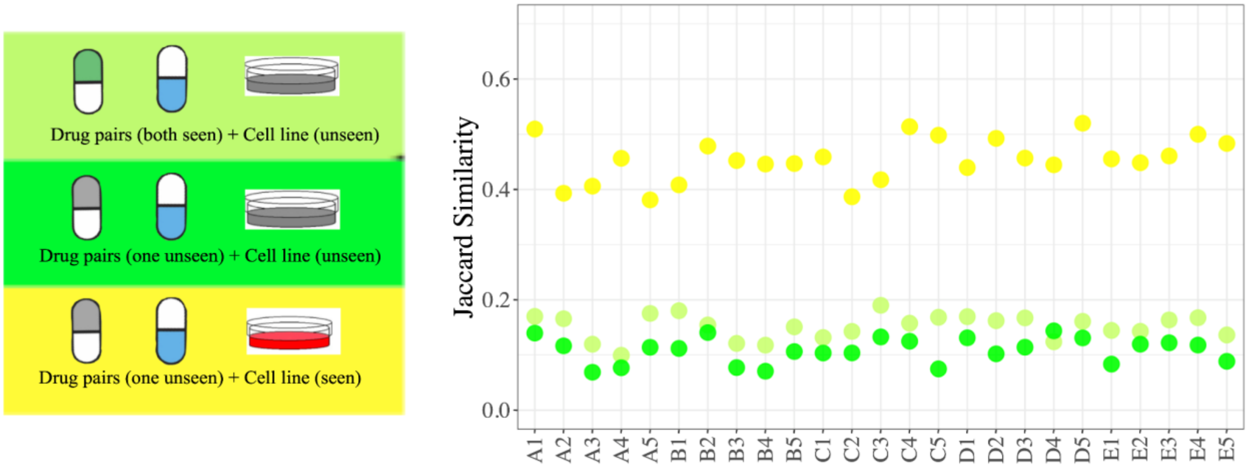


**Supplementary Figure S13. Agreement between CCSynergy III and V in three different triplet types across the 25 CC spaces.** Vertical axis indicates the Jaccard similarity (JS) between the set of synergistic triplets predicted by CCSynergy III with that of CCSynergy V in three scenarios described and color-coded at the legend (left) across the 25 CC spaces (horizontal axis). Note that in the seen-cell line scenario (yellow), in line with our expectations, the JS value across all CC spaces is considerably higher than the other two (un)seen-cell line scenarios.

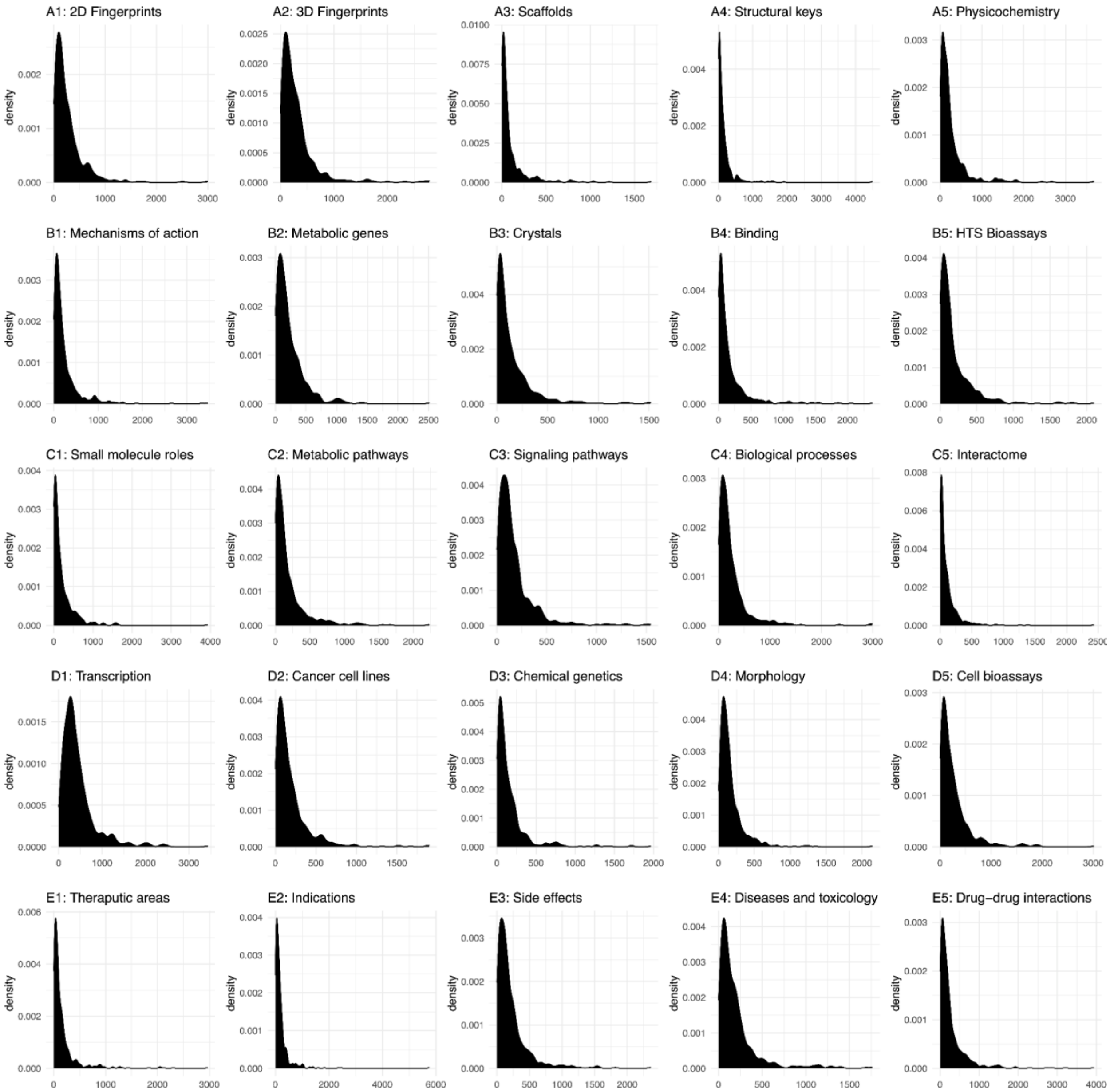


**Supplementary Figure S14.** Power-law distribution of the number of synergistic triplets per cell line (CCSynergy V) across the 25 CC spaces.

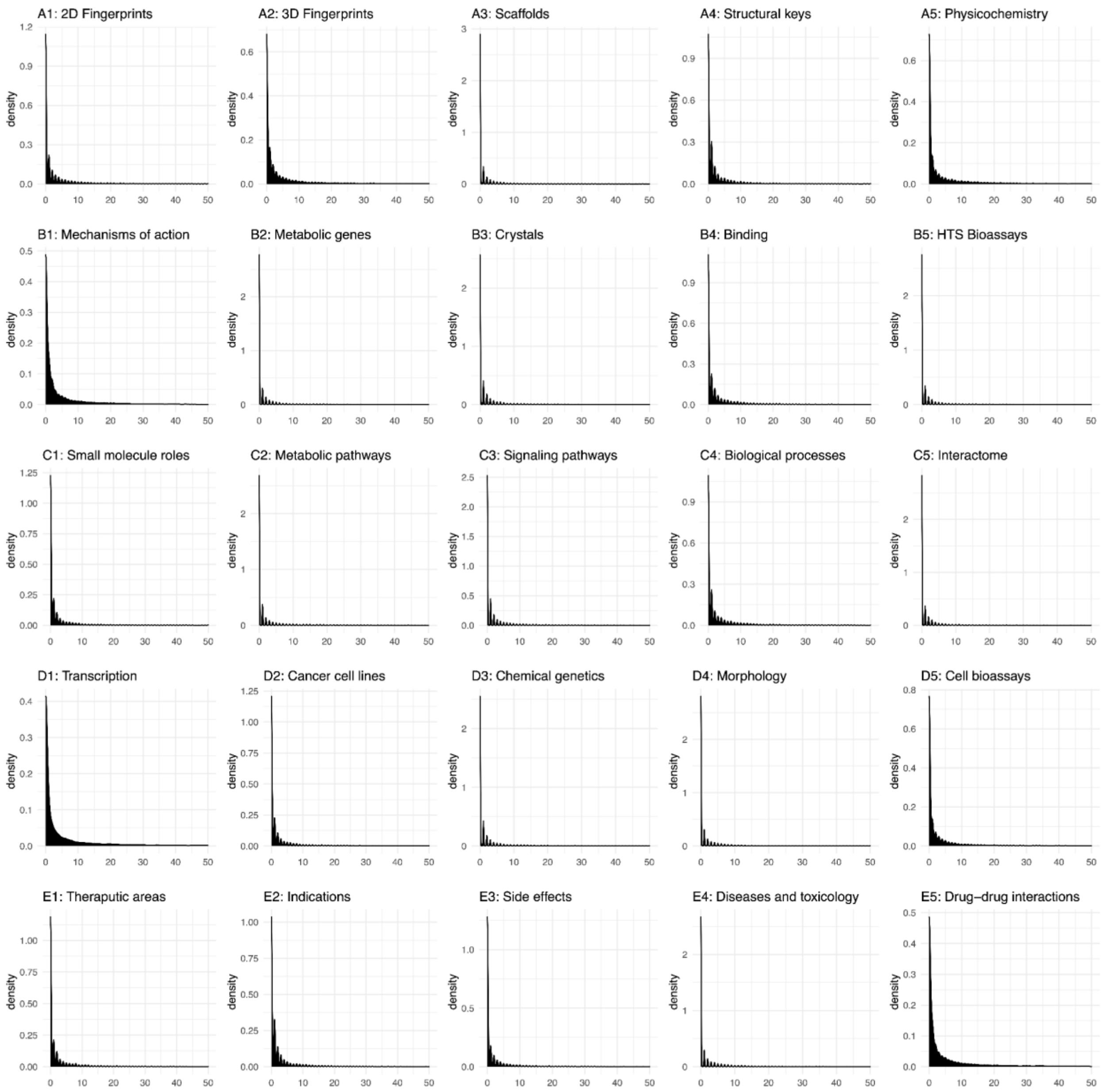


**Supplementary Figure S15.** Power-law distribution of the number of synergistic triplets per drug pair (CCSynergy V) across the 25 CC spaces.

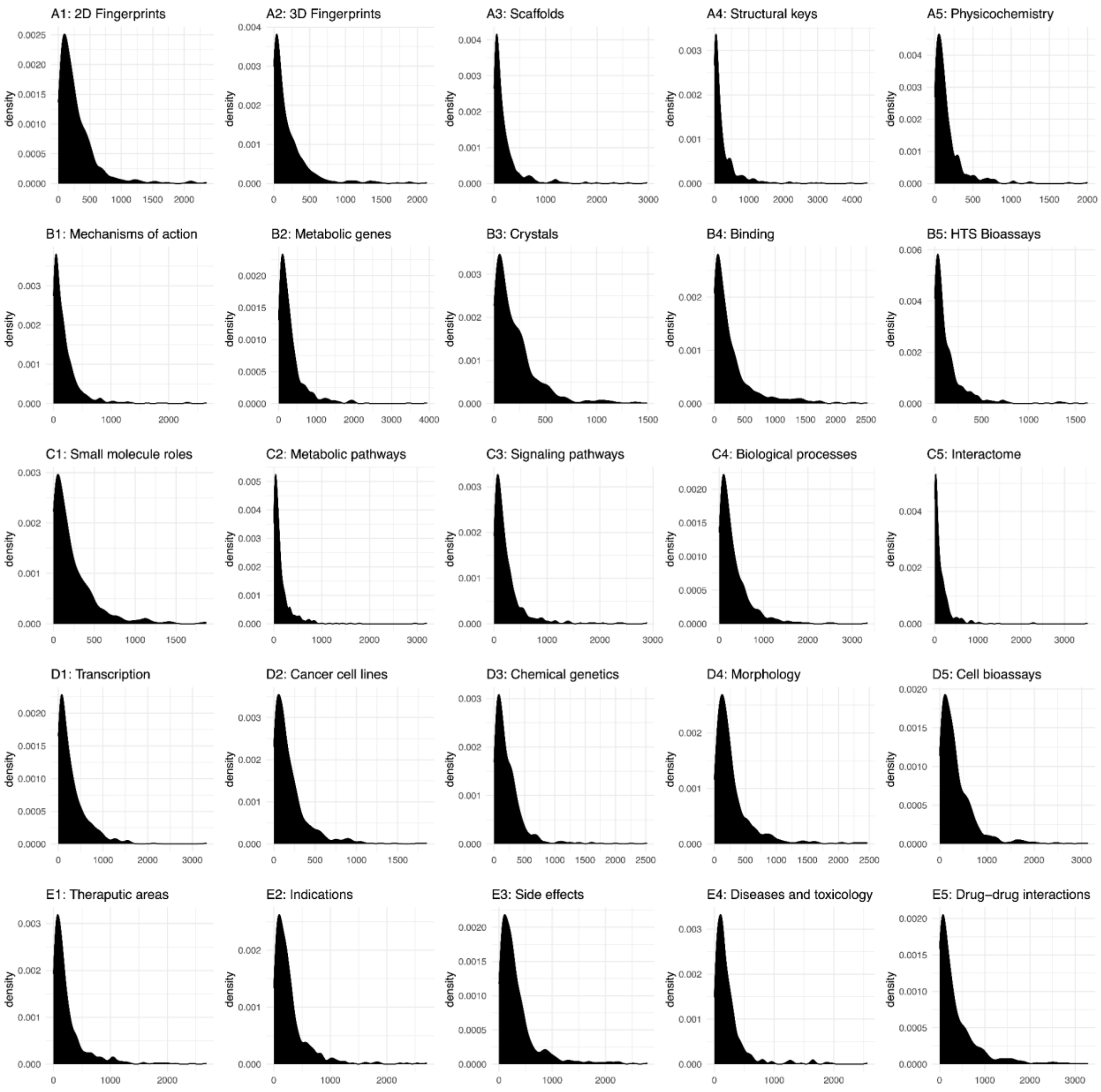


**Supplementary Figure S16.** Power-law distribution of the number of synergistic triplets per cell line (CCSynergy III) across the 25 CC spaces.

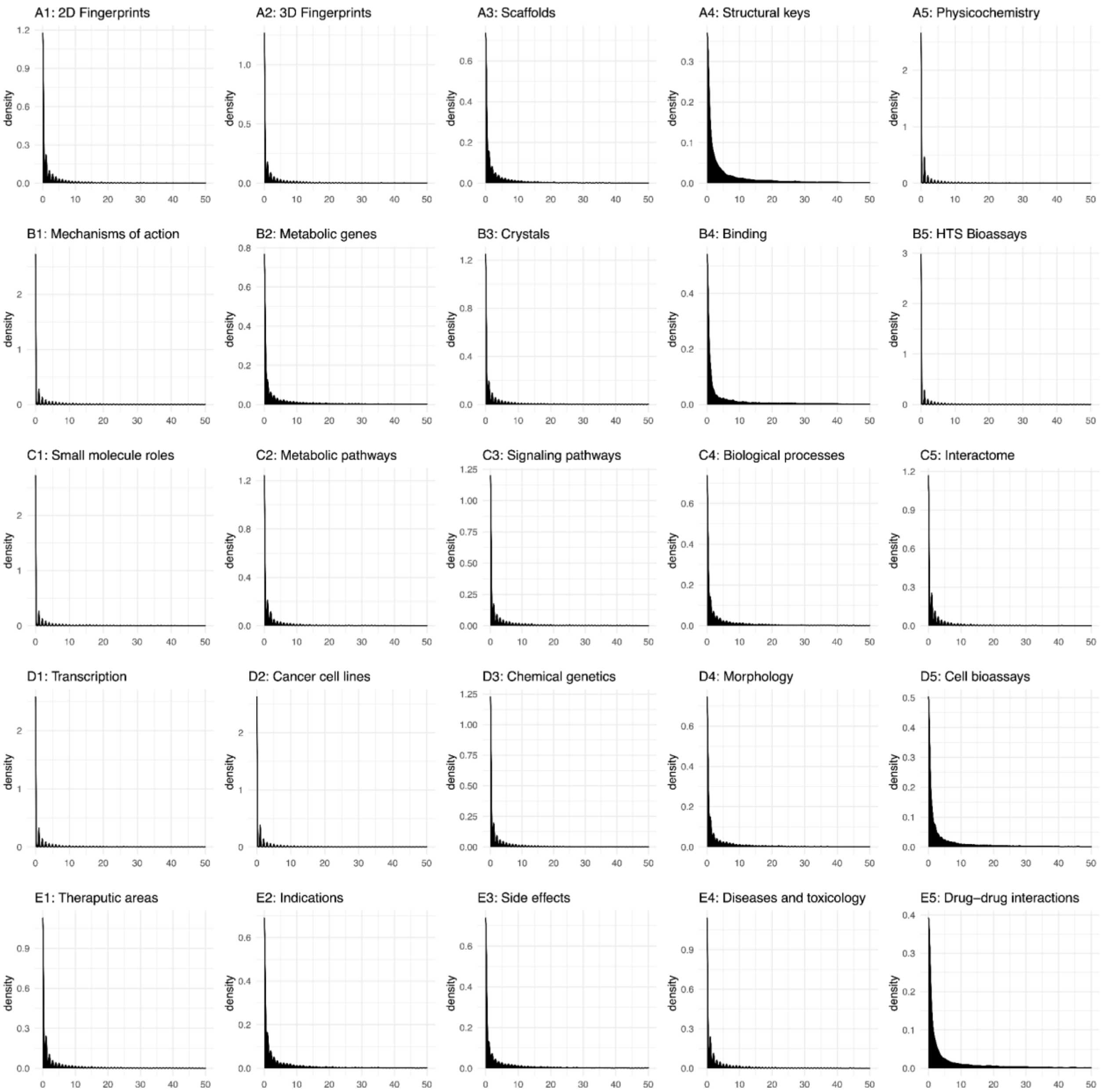


**Supplementary Figure S17.** Power-law distribution of the number of synergistic triplets per drug pair (CCSynergy III) across the 25 CC spaces.

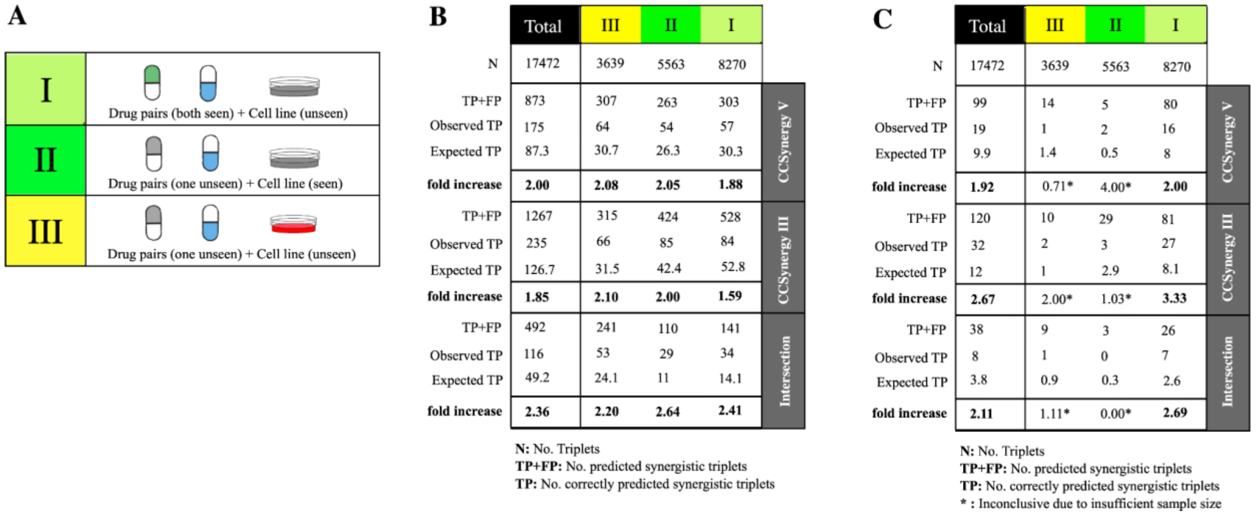


**B**

| | | Total | III | II | I |
|---|---|---|---|---|---|
| | N | 17472 | 3639 | 5563 | 8270 |
| CCSynergy V | TP+FP | 873 | 307 | 263 | 303 |
| CCSynergy V | Observed TP | 175 | 64 | 54 | 57 |
| CCSynergy V | Expected TP | 87.3 | 30.7 | 26.3 | 30.3 |
| CCSynergy V | **fold increase** | **2.00** | **2.08** | **2.05** | **1.88** |
| CCSynergy III | TP+FP | 1267 | 315 | 424 | 528 |
| CCSynergy III | Observed TP | 235 | 66 | 85 | 84 |
| CCSynergy III | Expected TP | 126.7 | 31.5 | 42.4 | 52.8 |
| CCSynergy III | **fold increase** | **1.85** | **2.10** | **2.00** | **1.59** |
| Intersection | TP+FP | 492 | 241 | 110 | 141 |
| Intersection | Observed TP | 116 | 53 | 29 | 34 |
| Intersection | Expected TP | 49.2 | 24.1 | 11 | 14.1 |
| Intersection | **fold increase** | **2.36** | **2.20** | **2.64** | **2.41** |

**N:** No. Triplets
**TP+FP:** No. predicted synergistic triplets
**TP:** No. correctly predicted synergistic triplets

**C**

| | | Total | III | II | I |
|---|---|---|---|---|---|
| | N | 17472 | 3639 | 5563 | 8270 |
| CCSynergy V | TP+FP | 99 | 14 | 5 | 80 |
| CCSynergy V | Observed TP | 19 | 1 | 2 | 16 |
| CCSynergy V | Expected TP | 9.9 | 1.4 | 0.5 | 8 |
| CCSynergy V | **fold increase** | **1.92** | 0.71* | 4.00* | **2.00** |
| CCSynergy III | TP+FP | 120 | 10 | 29 | 81 |
| CCSynergy III | Observed TP | 32 | 2 | 3 | 27 |
| CCSynergy III | Expected TP | 12 | 1 | 2.9 | 8.1 |
| CCSynergy III | **fold increase** | **2.67** | 2.00* | 1.03* | **3.33** |
| Intersection | TP+FP | 38 | 9 | 3 | 26 |
| Intersection | Observed TP | 8 | 1 | 0 | 7 |
| Intersection | Expected TP | 3.8 | 0.9 | 0.3 | 2.6 |
| Intersection | **fold increase** | **2.11** | 1.11* | 0.00* | **2.69** |

**N:** No. Triplets
**TP+FP:** No. predicted synergistic triplets
**TP:** No. correctly predicted synergistic triplets
*** :** Inconclusive due to insufficient sample size

**Supplementary Figure S18. Partial validation of CCSynergy (SML and MV-based integrations).** We identified partial overlap between the triplets considered in the CCSynergy database with those in the DrugComb database. We ranked triplets in this subset in terms of their Loewe synergy score, and considered the top 10% as synergistic (i.e., those with Loewe score > 9.2). **A)** We partitioned the triplets the same as in **Fig. 6b** and **Fig. 7a** in the main text. Note that the results in panels **B** and **C** were obtained based respectively on SML and MV integration of the 25 CC spaces. In both panels, the number of triplets in the overlapping set (N), number of synergistic triplets predicted by CCSynergy (TP+FP), number of triplets that in both databases are considered as synergistic (i.e., truly synergistic cases: observed TP), the TP that is expected by chance (10% of (TP+FP)), and the precision fold increase, which is basically the ratio of observed by expected TP, are shown for each three scenarios separately and in total (the column names). These measurements have been calculated both for CCSynergy V and III and also their intersection (the row names in the right-hand side). Note that in panel C, the results for scenarios II and III remain inconclusive due to insufficient synergistic sample size that is mainly a consequence of the stringency of the MV-based integration approach.

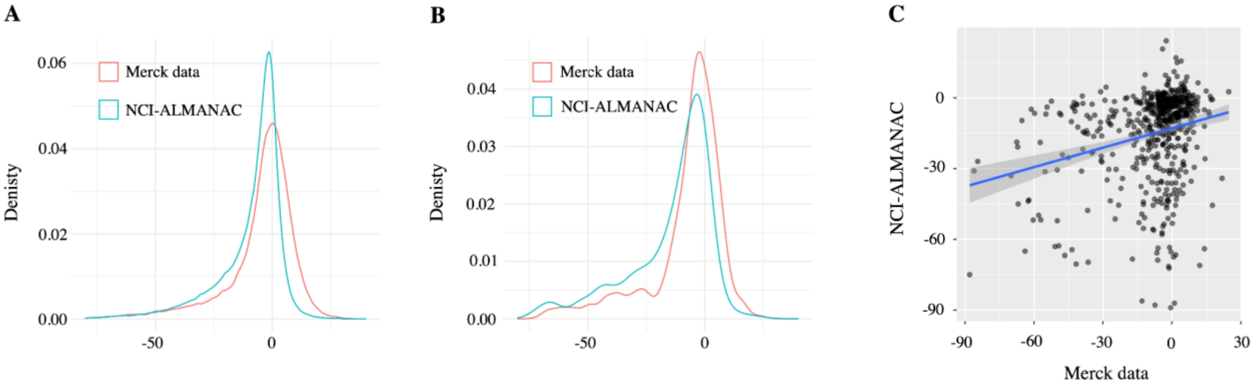


**Supplementary Figure S19. Poor reproducibility of Loewe scores across the Merck and NCI-ALMANAC datasets. A)** The overall distribution of Loewe scores in the Merck dataset (red curve) is shifted from that of NCI-ALMANAC (green curve). **B)** We identified 543 triplets that are shared between the two datasets and checked the distribution of the Loewe scores in this overlapping subset. We can see that the distribution shift still remains apparent. **C)** Scatter plot of the Loewe scores is shown, where points represent the triplets in the overlapping subset, and their *x* and *y* values indicate their corresponding Loewe scores in the Merck and NCI-ALMANAC datasets respectively. The blue line is the regression line and the Pearson Correlation Coefficient is 0.25, which is very low and confirms the poor reproducibility of synergy scores between different studies.
